# A simple regulatory network coordinates a bacterial stress response in space and time

**DOI:** 10.1101/2024.03.07.583862

**Authors:** Divya Choudhary, Kevin R. Foster, Stephan Uphoff

## Abstract

Bacteria employ diverse gene regulatory networks to protect themselves from stressful environments. While transcriptomics and proteomics show that the expression of different genes can shift strongly in response to stress, the underlying logic of large regulatory networks is difficult to understand from bulk measurements performed at discrete time points. As a result, it remains challenging to predict how these regulatory networks function at a system level. Here we use time-resolved single-cell imaging to explore the functioning of a key bacterial stress response: The *Escherichia coli* response to oxidative stress. Our work reveals a striking diversity in the expression dynamics of genes in the regulatory network, with differences in the timing, magnitude, and direction of expression changes. Nevertheless, we find that these patterns have a simple underlying logic. Firstly, all genes exhibit a transient increase in their protein levels simply due to the slowing down of cell growth under stress. Controlling for this effect reveals three classes of gene regulation driven by the transcription factor OxyR. Downregulated genes drop in expression level, while upregulated genes either show pulsatile expression that decays rapidly or gradual induction, dependent upon transcription factor binding dynamics. These classes appear to serve distinct functional roles in cell populations. Pulsatile genes are stress-sensitive and activate rapidly and transiently in a few cells, which provides an initial protection for cell groups. Gradually upregulated genes are less sensitive and induce more evenly generating a lasting protection that involves a larger number of cells. Our study shows how bacterial populations use simple regulatory principles to coordinate a stress response in space and time.

## Introduction

Bacterial stress response regulons are highly complex in that they commonly involve a suite of genes performing diverse functions, which display a variety of expression patterns under stress^1^. However, these regulons are also typically under the control of a single master transcriptional regulator. This begs the question of how such simple regulation is able to orchestrate the timing and magnitude of so many genes. Bulk transcriptomic and proteomic studies have documented an association between stress tolerance and gene regulation for a wide range of stresses, including under starvation^2,3^, oxidative stress^4^, osmotic stress^5^, heat stress ^5,6^, pH stress ^7^, DNA damage^8,9^, phage infection^10^, and antibiotic exposure^11^. Moreover, these methods often identify tens to hundreds of genes that shift in expression with stress treatments, which suggests a complex regulatory network underlying each response^12^. However, bulk population measurements are limited in their ability to link gene regulation with cellular phenotypes such as growth rate, because they lack single-cell resolution. In addition, due to cost and practicality, genome-wide gene expression data is typically recorded at only a few time points, meaning that temporal dynamics are not followed in any detail^13,14^. A final limitation of bulk measurements is an inability to capture phenotypic heterogeneity, whereby individual cells display different responses in space and time^15–20^.

An important alternative to bulk expression measurements is to use fluorescent reporter constructs that allow the gene expression in single cells to be followed. This approach has indicated the importance of temporal order in the activation of different genes within a regulon, e.g. for the SOS response, antibiotic stress, flagellar synthesis, and metabolic pathways ^8,9,16,18,21–24^. Spatial cell-cell variability can also emerge due to variability in the concentrations of stress agents and metabolites in cell populations^15,17,25^. The result can be complex spatio-temporal gene expression patterns under stress, which are shaped by cell-cell interactions, feedbacks between cell growth rate and gene expression^20,26,27^, and environmental fluctuations^19,20,28,29^. It is also becoming increasingly clear that stress survival is only partly determined by the protective responses of each individual cell, but strongly dependent on the collective protection provided by cell populations. However, to date, most single-cell studies have focussed on one or a few reporter genes for a given stress or disregarded the potential for cellular interactions, which leaves open the question of how whole stress response regulons are coordinated in cell populations.

Here, we leverage the recent advancements in single-cell imaging that allow cellular behaviour to be followed in a constantly controlled environment for long durations^18–20^. We focus on the oxidative stress response of *Escherichia coli* to hydrogen peroxide (H_2_O_2_). H_2_O_2_ is a major reactive oxygen species that causes damage to proteins, lipids, and DNA. H_2_O_2_ readily permeates cells^27^ where it causes redox imbalance ^30^. This stress leads to oxidation and a conformational change in the transcription factor OxyR^31,32^. OxyR acts as a sensor for intracellular H_2_O_2_ levels and controls an array of more than two dozen genes in its regulon, involved in diverse functions to maintain cell survival under H_2_O_2_ stress^4,33–36^, including metal ion homeostasis^37–41^, H_2_O_2_ scavenging^30,42,43^, and redox maintenance^44–46^. We combine theory and experiments to characterise and understand the transcriptional regulation of 31 genes, which have been previously identified as part of the oxidative stress response^4,30,32,36–42,44–60^ (Table 1). In this way, we are able to explain how a single transcription factor coordinates tens of genes in space and time to protect cells.

**Table 1:** Genes chosen for this study along with their characteristics.

| # | Gene | Function | Implicated in oxidative stress | Regulation in this study (peak /steady-state) | Promoter activity significance value (peak /steady-state) |
| --- | --- | --- | --- | --- | --- |
| 1 | *sodA* | Superoxide dismutase | P<sup>47</sup> | P/- | 1.72e<sup>-06</sup>/n.s. |
| 2 | *fhuF* | Ferric iron reductase | N<sup>4,48,49</sup> | N/N | 2.21e<sup>-18</sup>/2.46e<sup>-10</sup> |
| 3 | *trxC* | Thioredoxin 2 | P<sup>4,36,44-46,48,49</sup> | P/P | 2.83e<sup>-23</sup>/2.65e<sup>-24</sup> |
| 4 | *yaaA* | Suppress intracellular iron levels | P<sup>4,38,48</sup> | P/P | 2.23e<sup>-09</sup>/2.0e<sup>-12</sup> |
| 5 | *dps* | Fe-binding & storage | P<sup>4,36,49,51,53</sup> | N/- | 4.9e<sup>-05</sup>/n.s. |
| 6 | *ahpC* | Akylhydroperoxidase | P<sup>4,36,42,43,48,49</sup> | P/P | 1.45e<sup>-17</sup>/4.36e<sup>-06</sup> |
| 7 | *katG* | Catalase | P<sup>4,30,36,42,48,49</sup> | P/P | 5.24e<sup>-22</sup>/2.65e<sup>-24</sup> |
| 8 | *grxA* | Glutaredoxin-1 | P<sup>4,36,44,45,49</sup> | P/P | 7.029e<sup>-18</sup>/1.18e<sup>-18</sup> |
| 9 | *dsbG* | Sulfenic acid reductase | P<sup>4,49,54,79</sup> | N/- | 1.912e<sup>-09</sup>/n.s. |
| 10 | *znuA* | Zinc transport | P<sup>36</sup> | N/- | 7.61e<sup>-06</sup>/n.s. |
| 11 | *metE* | methionine synthase | P/N<sup>36,55</sup> | N/N | 3.73e<sup>-09</sup>/2.31e<sup>-06</sup> |
| 12 | *oxyR* | OxyR | N<sup>4,32,49</sup> | N/- | 1.512e<sup>-09</sup>/n.s. |
| 13 | *hemH* | Ferrochelatase | P<sup>4,41,48,49</sup> | P/P | 7.029e<sup>-18</sup>/1.18e<sup>-18</sup> |
| 14 | *mntR* | Mn<sup>2+</sup> regulator<sup>56</sup> | N<sup>4,49</sup> | N/N | 1.359e<sup>-10</sup>/0.015 |
| 15 | *gntP* | gluconate permease | N<sup>4,48,49</sup> | N/- | 1.512e<sup>-09</sup>/n.s. |
| 16 | *uxuA* | Mannonate hydrolase | N<sup>4,48,49</sup> | P/P | 3.36e<sup>-05</sup>/0.016 |
| 17 | *iscS* | Iron-sulfur cluster assembly | N<sup>40,48</sup> | N/- | 1.29e<sup>-09</sup>/n.s. |
| 18 | *hcp* | Hybrid cluster proteins of Fe/S | P<sup>4,49,57</sup> | -/- | n.s./ n.s. |
| 19 | *flu* | surface adhesin | N/P<sup>4,36,49,58</sup> | N/N | 0.0076/0.00545 |
| 20 | *ybjC* | Unknown | N<sup>4,36,48,49</sup> | N/N | 1.39e<sup>-09</sup>/0.00036 |
| 21 | *ychF* | ATPase | N<sup>4,49,59</sup> | N/- | 2.77e<sup>-06</sup>/n.s. |
| 22 | *yfdL* | Putative ligase | -<sup>79</sup> | -/- | n.s./ n.s. |
| 23 | *metR* | methionine biosynthesis<sup>80</sup> | P<sup>4,36</sup> | N/N | 6.046e<sup>-09</sup>/3.73e<sup>-06</sup> |
| 24 | *znuC* | Zinc transport<sup>81</sup> | P<sup>4,36</sup> | N/- | 1.34e<sup>-06</sup>/n.s. |
| 25 | *ybjN* | Unknown | N<sup>4,49</sup> | -/- | n.s./ n.s. |
| 26 | *elaB* | Membrane integrity | P<sup>4,60</sup> | P/P | 2.05e<sup>-05</sup>/n.s. |
| 27 | *fur* | iron-homeostatic control protein | P<sup>4,39,48,49</sup> | P/P | 2.44e<sup>-12</sup>/5.60e<sup>-13</sup> |
| 28 | *poxB* | pyruvate:quinoneoxidoreductase<sup>82</sup> | P<sup>4,48</sup> | P/P | 8.58e<sup>-07</sup>/6.17e<sup>-05</sup> |
| 29 | *yaiA* | Unknown | P<sup>4,48</sup> | P/P | 0.00015/0.00279 |
| 30 | *clpS* | Iron sequestration | P<sup>37</sup> | P/P | 1.3877e<sup>-13</sup>/1.18e<sup>-18</sup> |
| 31 | *clpX* | Iron sequestration | P<sup>37</sup> | N/N | 1.738e<sup>-05</sup>/3.67e<sup>-06</sup> |

P: upregulation, N: downregulation; p-values using Mann-Whitney tests where n.s. for p>0.05

## Results

### Genes in the OxyR regulon show variable responses to stress

We imaged gene expression dynamics using time-lapse microscopy of *E. coli* cells, each carrying one of 31 plasmid-based transcriptional reporters that have an OxyR-regulated promoter followed by a fast-maturing GFP^61^ or sCFP3 (for P*katG*, P*ahpC* and P*grxA*^28,62^). We collected data for ∼10,000 individual cells growing inside microfluidic growth trenches (‘mother machine type’) during continuous H_2_O_2_ stress from ∼2 hours before until 6 hours of treatment at 3-min time intervals, resulting in a rich dataset of ∼200,000 expression measurements per reporter gene [Figure 1A]. The microfluidic chip provides an opportunity to disentangle genuine gene regulatory dynamics from environmental changes because the continuous inflow of fresh media with H_2_O_2_ keeps the treatment concentration constant despite changes in the H_2_O_2_ scavenging capacities of cells [Figure 1B]. Another feature of our approach is to study the AB1157 strain of *E. coli* that carries an amber mutation in RpoS^63^. This allows us to isolate the role of OxyR in gene regulation without the influence of stationary phase regulation of certain oxidative stress response genes by RpoS^64–66^.

**Figure 1:**
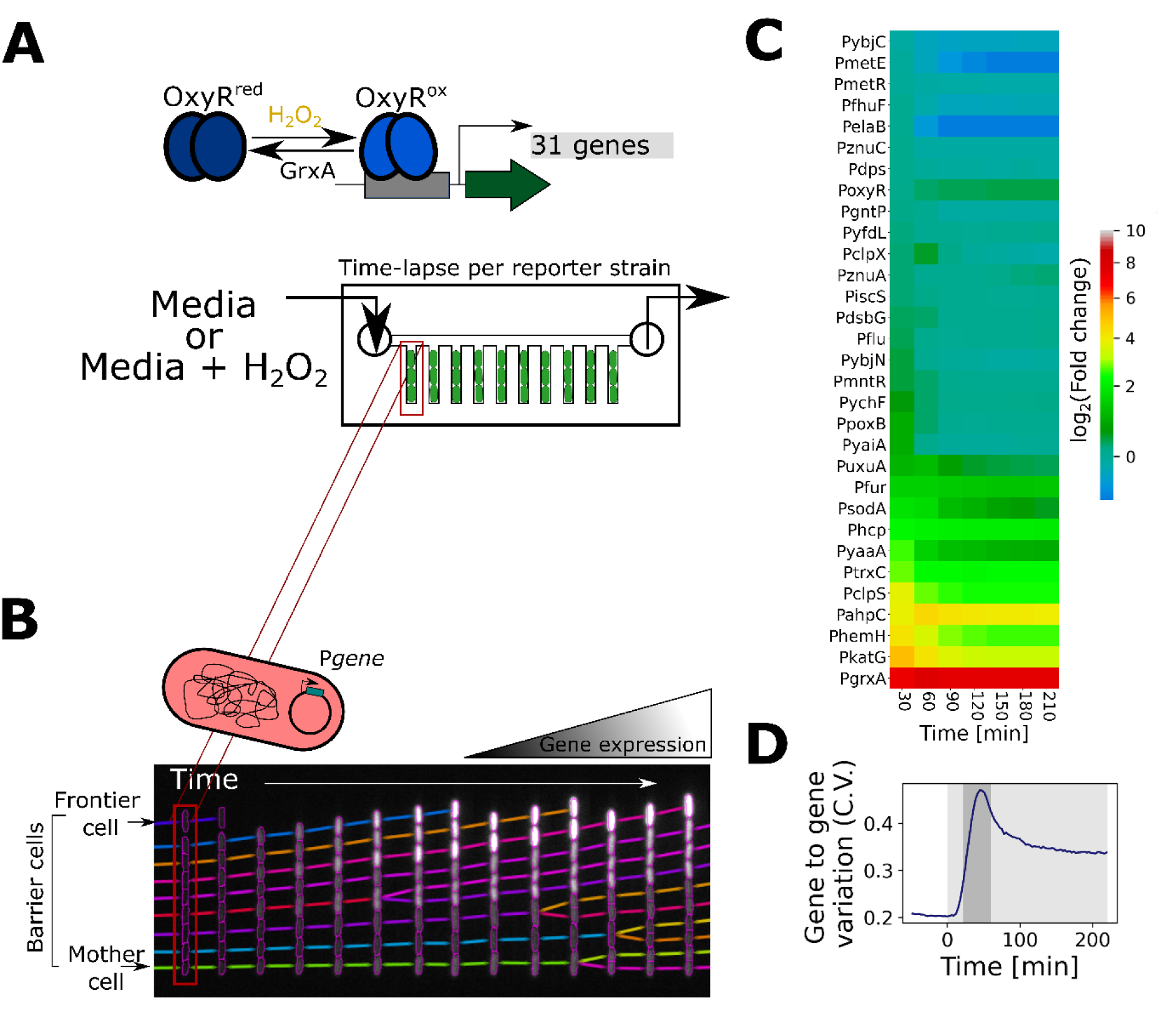
Gene expression dynamics across the OxyR regulon under H_2_O_2_ treatment. (**A**) Upon exposure to hydrogen peroxide, oxidation of OxyR leads to induction of a wide regulon of genes in *E. coli*. The schematic displays the microfluidics-based methodology to visualise the promoter activity of 31 fluorescently-tagged transcriptional reporters by time-lapse microscopy at single-cell resolution. (**B**) The kymograph represents cell growth over time in one of the representative growth channels with intensity of cells reporting gene expression levels. We define ‘mother cells’, ‘barrier cells’, and ‘frontier cells’ according to their position relative to the open end of the growth channel where growth media with or without H_2_O_2_ treatment is provided. (**C**) Heatmap represents mean log_2_-fold change in gene expression relative to basal level of frontier cells for 31 transcriptional reporters at 30-minute intervals during continuous treatment with 100 µM H_2_O_2_ from t = 0 min (n ≥ 1000 cells and ≥2 repeats per gene). (**D**) Coefficient of variation (C.V.) for gene-to-gene expression variability across the 31 transcriptional reporters of frontier cells under 100 µM H_2_O_2_ treatment provided at t = 0 minutes. (Shaded region represents H_2_O_2_ treatment, with the darker shade representing the highly variable gene-gene expression post treatment; n ≥ 1000 cells and ≥ 2 repeats per gene).

With this setup, we were able to observe that all 31 genes showed an initial expression pulse within 10 min after the start of treatment [Figure 1C], but expression of different genes diverged after this initial pulse. This variability fits with observations on the expression dynamics of the oxidative stress response of *Pseudomonas aeruginosa*^67^. After 90 min of continuous H_2_O_2_ treatment, most genes remained upregulated while others became downregulated relative to the untreated expression level [Figure 1C]. To further characterise differences in the regulation between genes, we computed the coefficient of variation (CV, variance normalised by the mean) across the expression levels of the 31 genes. The gene-to-gene variation was low in untreated cells, greatest during the transient expression peak shortly after treatment, and intermediate during steady-state with continuous H_2_O_2_ treatment [Figure 1D]. These patterns suggest that the genes vary not only in expression magnitude but also show pronounced differences in induction timing and dynamics.

### Transient growth inhibition causes a genome-wide expression pulse during stress

Our methodology enables us to analyse individual cell growth dynamics alongside gene expression. Importantly, this analysis revealed that the expression pulse at the start of H_2_O_2_ treatment precisely coincides with a period of transient growth inhibition [Figure 2A]. During normal growth, protein concentrations stay constant in a cell because the expression rate is balanced with the dilution of molecules due to cell growth and division. H_2_O_2_ stress is known to rapidly stall cell growth which is triggered by an OxyS-mediated cell cycle arrest^68^. To remove the effect of growth dynamics on genes in the OxyR regulon, we normalised changes in expression level by the cell growth rate, to give a measure of the promoter activity^16^. Correcting for growth rate effects revealed that 3 of the 31 genes (P*hcp*, P*yfdL*, P*ybjN*) showed no significant change in promoter activity at any time [Figure 2B, C, Movie S1-S2]. Moreover, another 9 of the genes (7 downregulated and 2 upregulated) showed only a transient change in regulation during the first ∼100 minutes of treatment. These loci return to a baseline expression levels even though H_2_O_2_ is still present at a constant external concentration. Focusing our characterisation on genes with lasting expression changes left 11 upregulated genes and 8 downregulated genes that showed a prolonged response to stress.

**Figure 2:**
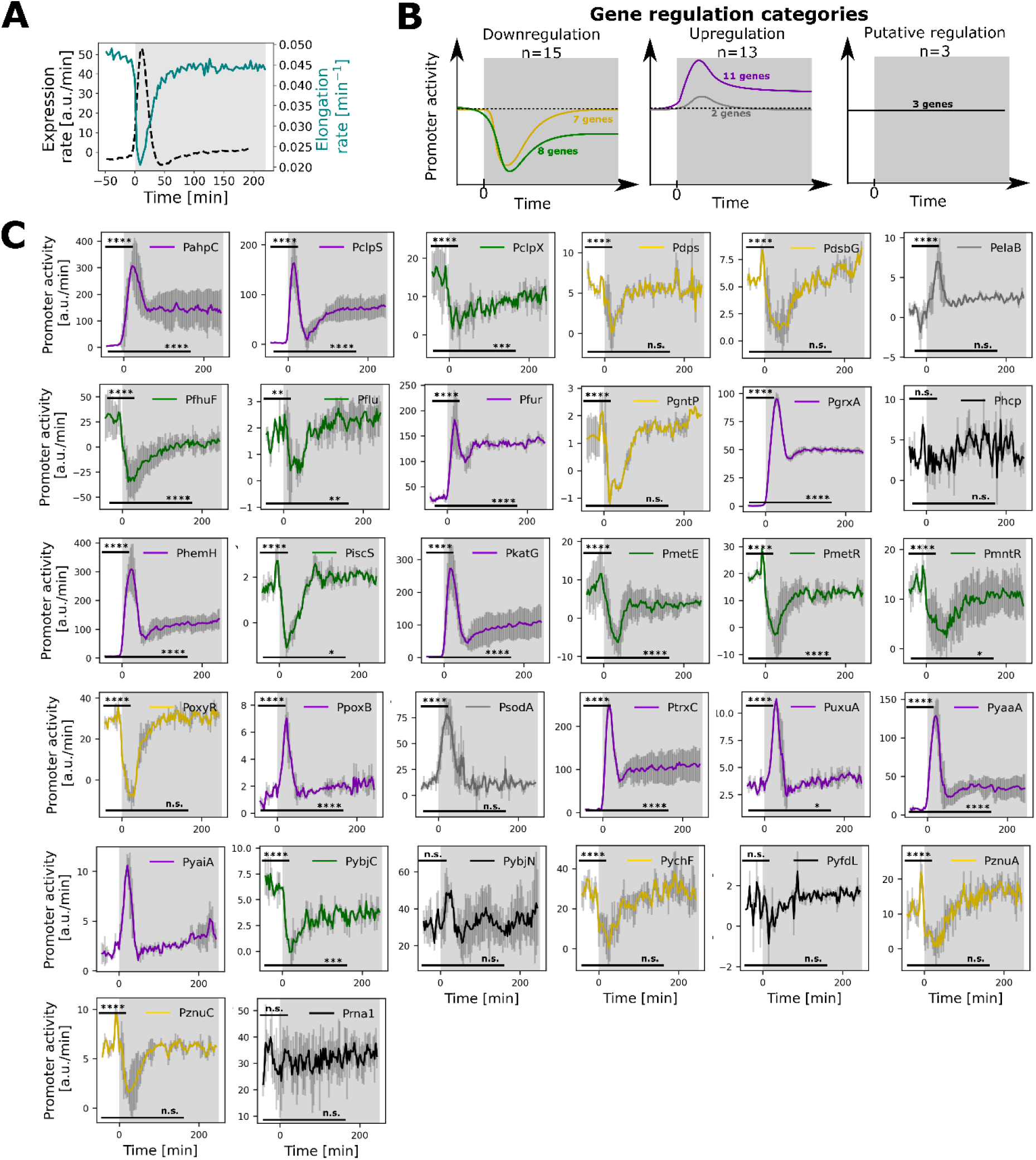
Oxidative stress response genes show a range of different expression dynamics during constant H_2_O_2_ stress. (**A**) Mean elongation rate (teal) and mean expression rate for all genes (black; dashed) of frontier cells treated with 100 μM H_2_O_2_ from t=0 min (shaded region). (**B**) Schematic illustrates the expression dynamics of the 5 gene regulation categories under H_2_O_2_ treatment (from t=0, shaded region) together with the number of genes in each category: (left) downregulation throughout the H_2_O_2_ treatment (green) or transient downregulation post treatment (gold); (middle) upregulation throughout the H_2_O_2_ treatment (purple) or transient upregulation post treatment (grey); (right) no significant change in promoter activity observed (black). Dashed lines represent the basal level of expression before treatment. (**C**) Mean promoter activity of frontier cells for 31 transcriptional reporters (ordered alphabetically) and the constitutively expressed promoter P*rna1* with constant 100 µM H_2_O_2_ treatment from t = 0 minutes (shaded region). Promoter activity shows the expression rate corrected for changes in cell growth rate. Non-parametric Mann-Whitney tests were used to indicate significant changes in promoter activity at the transient or steady-state relative to the basal level expression; where ****: p≤0.0001, ***: p≤0.001, **: p≤0.01, *: p≤0.05, and n.s.:p>0.05. Line colours correspond to the gene regulation categories as explained in panel B. Error bars represent standard deviation (n ≥ 1000 cells and ≥ 2 repeats per gene).

An interesting consequence of the effects of growth arrest on protein levels in the cell is that the genes that we identify as negatively regulated still show a positive pulse in protein levels after treatment [Figure 2B, C]. In other words, the transcription rate from these promoters is downregulated in response to H_2_O_2_, but the reduction in growth rate outweighs this effect, resulting in an effective accumulation of proteins and the illusion of gene upregulation. These conclusions are supported by studying the constitutively-expressed P*rna1* promoter that is not part of the OxyR regulon, which our analysis shows has a similar expression pulse but constant promoter activity [Figure 2C, Movie S3]. That is, we find that the growth arrest-dependent accumulation of proteins is a global genome-wide effect and not specific to oxidative stress response genes.

### Upregulated genes display either pulsatile or gradual dynamics

We next focused on the 11 genes whose gene expression changes and remains upregulated with H_2_O_2_ treatment [Figure 2B, S1]. Because we had found that gene-to-gene differences in expression were largest during the transient expression peak, we plotted the reporter expression level during the transient peak versus the level at steady-state [Figure 3D]. This analysis reveals that the upregulated genes cluster on 2 distinct slopes. Seven of the upregulated genes (P*katG*, P*yaaA*, P*clpS*, P*hemH*, P*uxuA*, P*poxB*, P*yaiA)* all have a high ratio of *peak/steady-state* ∼ 3, while the other upregulated genes (P*grxA,* P*trxC,* P*fur*, P*ahpC*) share a lower ratio of *peak/steady-state* ∼ 1.5 [Figure 3E]. The first class of genes showed a strong initial expression pulse that decays rapidly after its peak and stabilises at a low steady-state level (hence termed “pulsatile”). In contrast, the second class of genes with lower *peak/steady-state* ratio showed a gradual induction to an elevated steady-state level (hence termed “gradually induced”) [Figure 3B, C, Movie S1-S2]. After classifying the up-regulated genes into two groups based on the *peak/steady-state* ratio, we also found that the pulsatile genes are generally activated earlier after the onset of H_2_O_2_ stress, achieving their maximal expression in 25.7±4.7 minutes compared to 37.5±7.8 minutes post H_2_O_2_ treatment for the non-pulsatile genes [Figures 3F, S1, Movie S1-S2].

**Figure 3.**
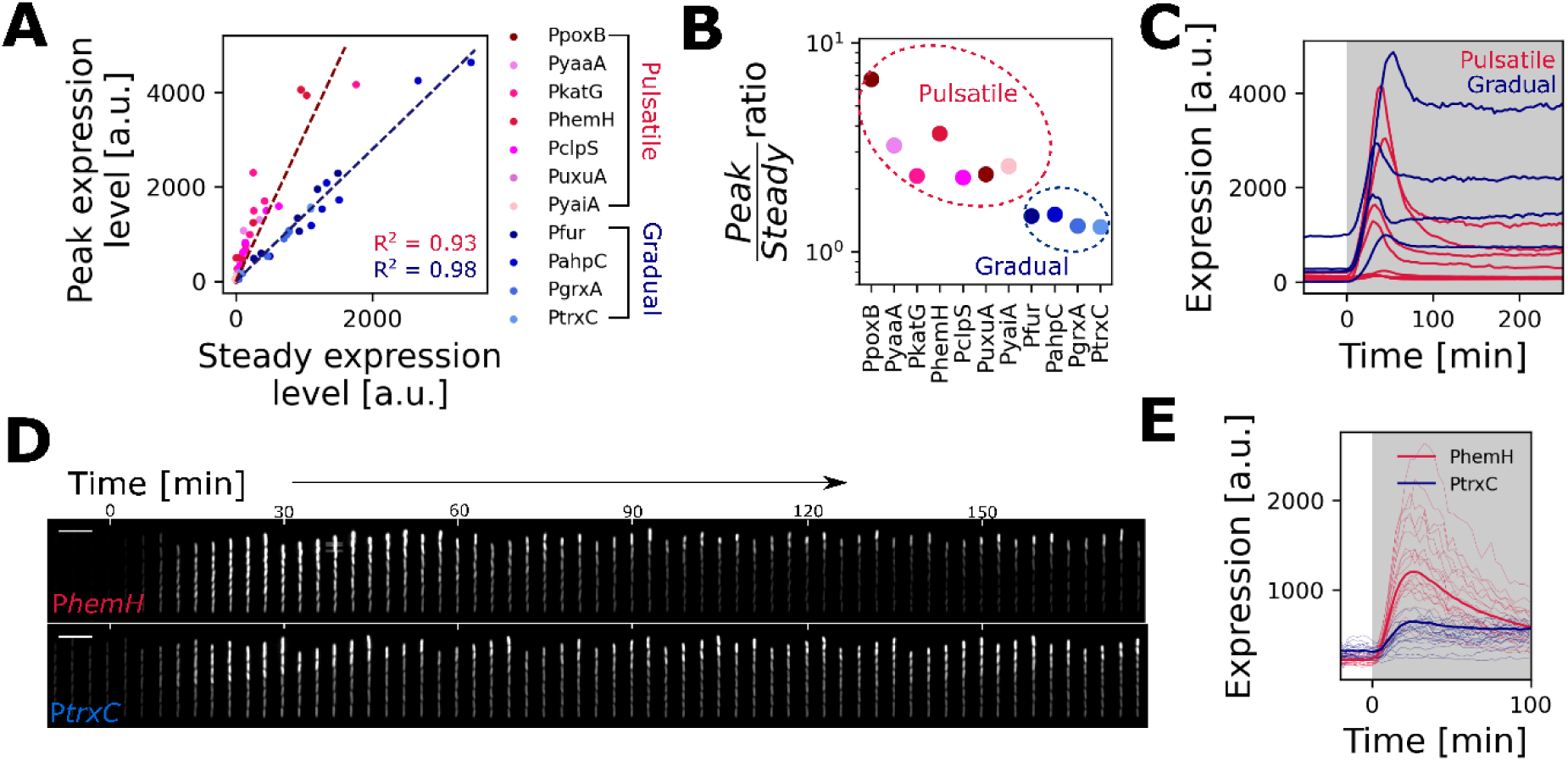
Upregulated promoters show either pulsatile or gradual induction dynamics. (**A**) Plot of the peak expression level (during the transient phase) versus the expression level at steady-state of frontier cells relative to basal expression for the indicated transcriptional reporters with 25 µM, 50 µM and 100 µM H_2_O_2_ treatment. The pulsatile (pink) and gradually induced (blue) genes cluster on two distinct slopes, shown with linear fits (dashed lines). (**B**) Peak/steady ratio obtained by linear regression of values for individual genes shown in panel A cluster into two categories (pink: pulsatile genes, blue: gradual induced genes, n≥3 repeats per gene). (**C**) Plot shows mean reporter expression levels of frontier cells for pulsatile (pink) and gradual (blue) genes under 100 µM H_2_O_2_ treatment provided at t = 0 min (shaded region) (n ≥ 1500 and n≥3 repeats per gene). (**D**) Kymograph of P*hemH* and P*trxC* transcription reporter expression representative of pulsatile and gradual induction with 100 µM H_2_O_2_ from t=0 minutes, respectively (Scale bar = 10 µm). (**E**) Plot shows mean (bold) and single cell (thin) reporter expression levels of mother cells for P*hemH* (pink, pulsatile) and P*trxC* (blue, gradual) under 100 µM H_2_O_2_ treatment provided at t = 0 min (shaded region).

### Modelling explains how differences in OxyR binding generate pulsatile or gradual dynamics

What explains the different patterns of upregulation? The oxidative stress response involves feedbacks from various metabolic pathways ^4,32,36,41,48,49,53–55^ and several genes in the OxyR regulon are co-regulated by more than one transcription factor^36,64–66,69,70^. Differences in the expression of genes in the OxyR regulon, therefore, may arise in multiple ways. However, we hypothesised that the differences we are seeing in the two classes of upregulated genes has a simpler underlying cause in the properties of OxyR itself. To explore this, we turned to a published minimal mathematical model from our previous work, which uses ordinary differential equations to describe changes in gene expression over time. Here, H_2_O_2_ permeates through the cell membrane and oxidises OxyR, which leads to the induction of genes of the oxidative stress response regulon^71^ [Figure 4A]. For simplicity, we reduced the operon to include only *grxA* (glutaredoxin-1) that converts OxyR back to its reduced form, the genes *ahpCF* (alkyl hydroperoxidase) and *katG* (catalase) that scavenge intracellular H_2_O_2_, and a ‘reporter gene’ that is induced by OxyR but does not have a functional role (to mimic the GFP reporters used in experiments). Gene expression increases on induction by OxyR and decreases due to dilution by cell growth. The growth rate slows down with increasing H_2_O_2_ concentration.

**Figure 4.**
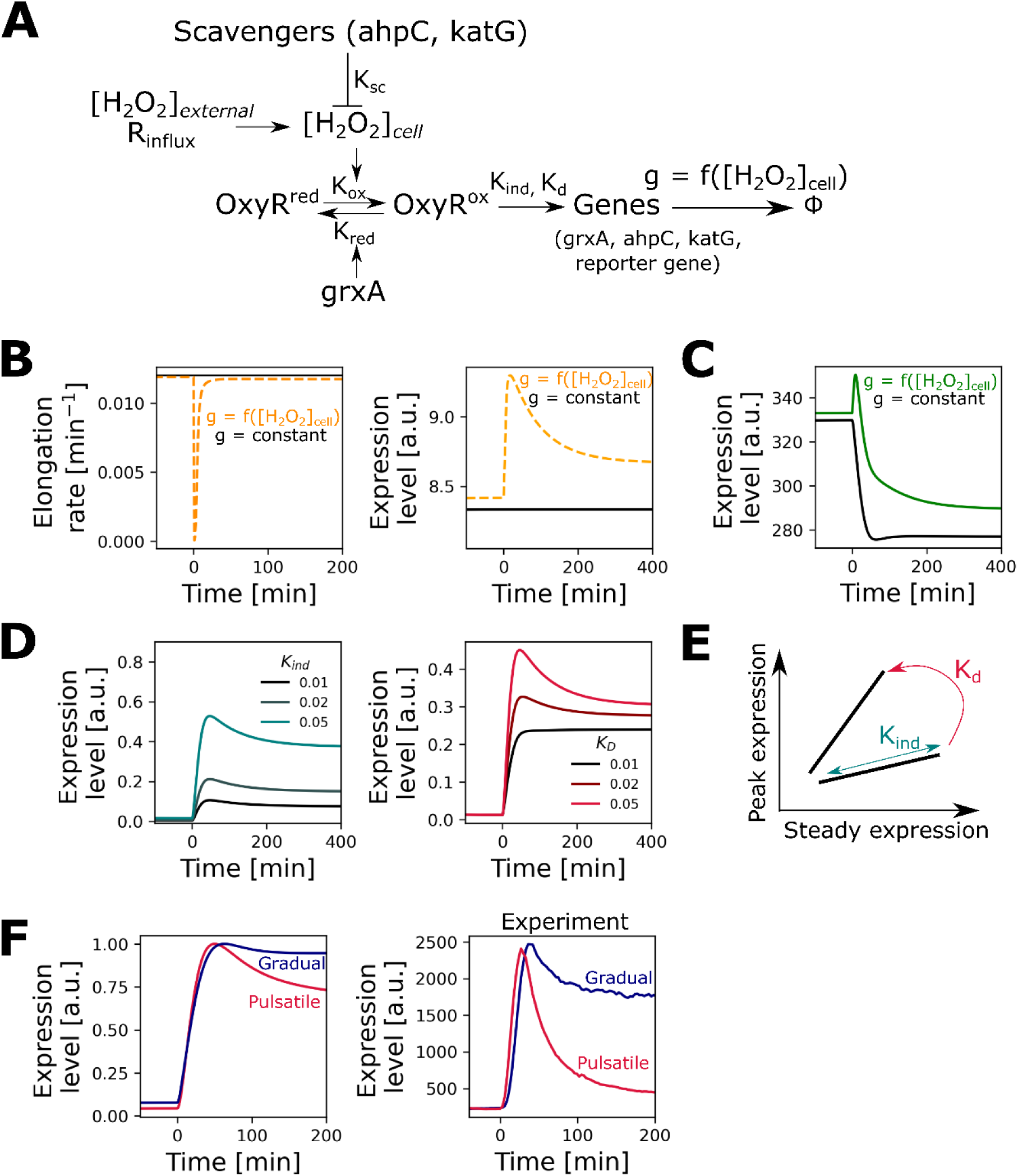
A model of the oxidative stress response predicts the molecular basis of the different categories of gene regulation. (**A**) Schematic represents the oxidative stress response model. The cells experience influx of H_2_O_2_ at R_influx_ rate, causing OxyR oxidation and induction of stress response genes. GrxA (glutaredoxin-1) converts oxidised OxyR back to its reduced form and scavenging enzymes AhpC and KatG lower the intracellular H_2_O_2cell_ concentration. The expression of the proteins is balanced by dilution due to cell growth, where the cell elongation rate g is a function of H_2_O_2cell_. OxyR also regulates a reporter gene which has no function in the response itself. (**B**) Effect of growth inhibition by H_2_O_2_ on gene expression dynamics. (left) Cell elongation rate affected by intracellular H_2_O_2_ concentration (orange, dashed) compared to constant growth rate unaffected (black) by H_2_O_2_ treatment provided at t = 0 min. (right) Expression level of a constitutively-expressed reporter gene (not regulated by OxyR) shows a passive induction pulse in the case where H_2_O_2_ inhibits cell elongation (orange) and no induction effect when the elongation rate is unaffected by H_2_O_2_ (black) treatment from t=0 min. (**C**) Expression dynamics of a downregulated reporter gene in the case where cell elongation rate is affected (green) or unaffected (black) by H_2_O_2_. (**D**) Induction of upregulated reporter genes for varying values of induction rate (*K_ind_*, left) and promoter dissociation constant of oxidised OxyR (*K_D_*, right) with H_2_O_2_ treatment from t = 0 min. *K_ind_*/*K_D_* ratio was kept constant for curves with varying *K_D_*. (**E**) Schematic represents the effect of *K_ind_* (teal) and *K_D_* (pink) on the gene expression dynamics. Pulsatile genes with high *peak/steady ratio* are characterised by a higher *K_D_* value, while *K_ind_* determines the overall magnitude of the induction without affecting the *peak/steady ratio*. (**F**) Mean gene expression levels from model (left) and experiments (right) of representative genes for the two categories of upregulation – P*katG* (pink, pulsatile gene) and P*ahpC* (blue, gradually induced gene) of frontier cells treated with 100 µM treatment from t = 0 min (n≥3 repeats).

We previously showed that this model recapitulates many aspects of the gene regulatory response and growth dynamics of *E. coli* with H_2_O_2_ treatment^71^. Here, we employ the model to understand the basis for the different expression patterns seen across the genes in the OxyR regulon. We first tested the model’s ability to explain the passive gene induction caused by the growth arrest-dependent accumulation of proteins. As predicted, when the expression of the reporter gene was uncoupled from OxyR regulation in the model, genes still show the expression pulse during the transient growth arrest while the promoter activity stayed constant [Figure 4B]. In addition, for a reporter gene that is downregulated by OxyR, we again observe a similar induction pulse followed by a decrease in expression [Figure 4C].

To further explain the data, we extended the model to explicitly describe the effects of OxyR-dependent gene regulation based upon two parameters: *K_ind_* captures the maximal expression rate when the promoter is fully occupied by oxidised OxyR, and *K_D_* captures the dissociation constant of oxidised OxyR from the promoter. Varying only these two parameters allowed us to shift expression dynamics in upregulated genes from gradual to pulsatile as observed in the data. Specifically, changing *K_D_* affects the *peak/steady-state* ratio of the gene, with pulsatile genes being characterised by a high *K_D_* value [Figure 4D]. *K_ind_* meanwhile determines the position of the gene along the slope in a *peak vs steady-state* plot. Our model predicts, therefore, that the two categories of gene regulation, pulsatile and gradual are the result of distinct promoter dissociation constants of OxyR [Figure 4D, E]. In support of this, experiments showed that the dissociation constant of oxidised OxyR from the pulsatile gene promoter P*katG* is an order of magnitude higher than for the gradually induced P*ahpC* promoter^72^ [Figure 4F]. We conclude, therefore, that the divergent patterns in gene regulation in the regulon can be explained by coupling a simple OxyR redox switch with patterns of cell growth under stress.

### Pulsatile genes protect against sudden stress

What is the benefit of pulsatile expression of some genes and gradual induction of others? The model shows how the dissociation constant of OxyR affects sensitivity of genes to changes in H_2_O_2_ concentration. Specifically, promoters with a high *K_D_* for OxyR show an approximately proportional increase in activity with H_2_O_2_ concentration, whereas the activity of promoters with a low *K_D_* is less sensitive to H_2_O_2_ [Figure 5A]. In support of this prediction, experiments show that rising H_2_O_2_ concentration causes a much steeper increase for the pulsatile P*katG* promoter activity compared to the gradually induced P*grxA* promoter [Figure 5A, S2]. Similarly, other pulsatile genes showed a higher sensitivity to H_2_O_2_ compared to gradually induced genes [Figure 5B, S2]. In *Mycobacterium tuberculosis*, induction of *katG* showed higher sensitivity for H_2_O_2_ than *ahpC* ^73^, matching the pulsatile behaviour of *katG* and gradual induction of *ahpC* in *E. coli*. The model further predicts that the pulsatile gene induction is a consequence of a response to a transient spike in intracellular H_2_O_2_ which accumulates until the increased level of scavenging enzymes tips the balance [Figure 5C]. In support of this explanation, experiments show that the expression pulse of P*katG* is much reduced when cells respond to a gradually increasing dose of H_2_O_2_ as opposed to a step treatment [Figure 5D].

**Figure 5.**
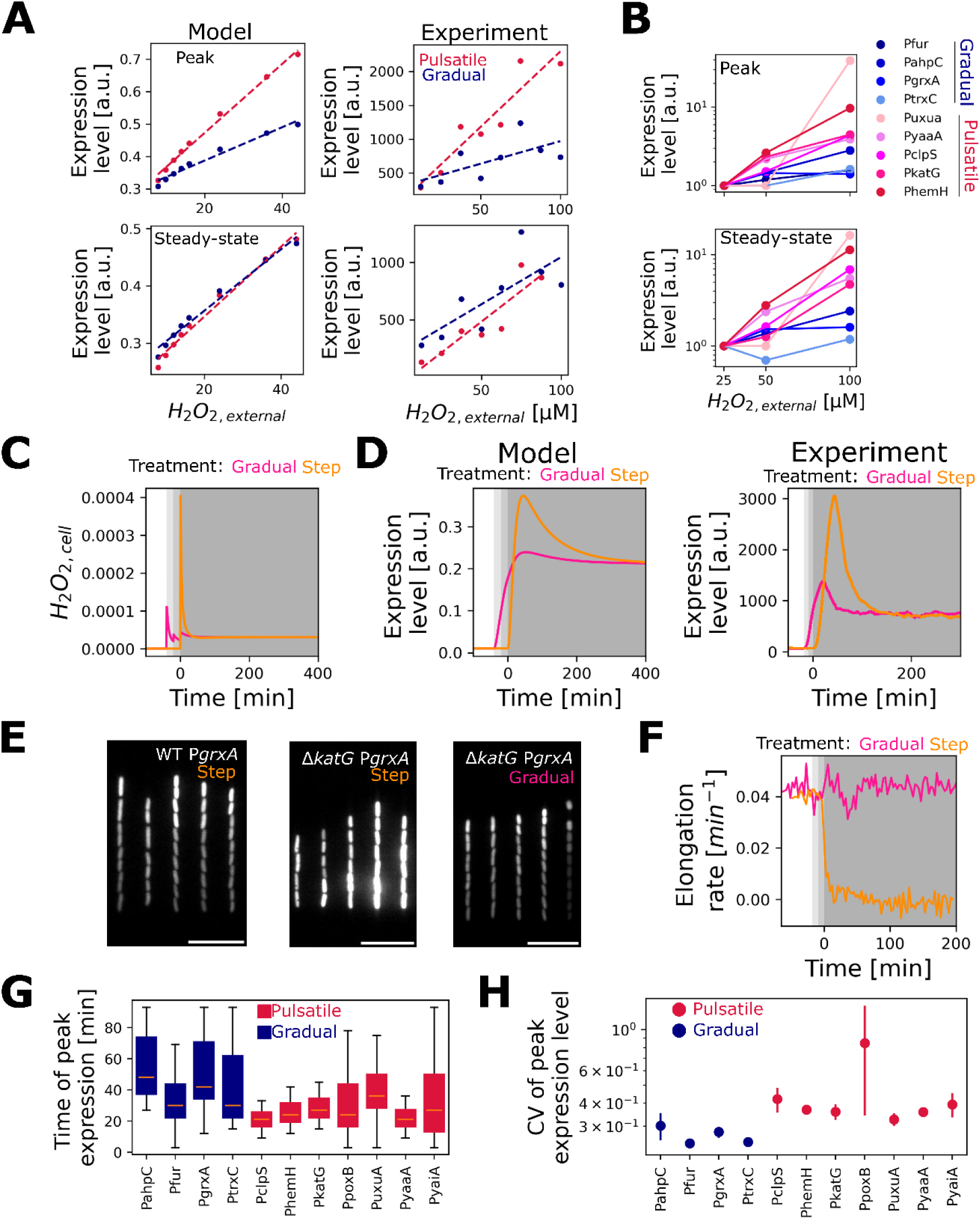
Pulsatile genes respond quickly to bridge the adaptation lag after H_2_O_2_ treatment. (**A**) The response of pulsatile genes is more sensitive to changes in H_2_O_2_ concentration than gradually induced genes. Model outputs (left) and experimental data (right) for expression level at peak (bottom) and steady state (bottom) of P*katG* (pulsatile gene, pink) and P*grxA* (gradual, blue) across a range of external H_2_O_2_ concentrations. Dashed lines show linear fits. (**B**) Relative changes in gene expression at peak (top) and steady-state (bottom) of frontier cells with H_2_O_2_ treatment ranging from 25 µM to 100 µM H_2_O_2_. Pulsatile genes (pink) show higher dose-sensitivity compared to gradual induced genes (blue) (n≥2 repeats per gene per H_2_O_2_ concentration). (**C**) Model results of intracellular H_2_O_2cell_ concentration with step treatment (100 µM H_2_O_2_, orange) from t= 0 minutes or gradual treatment (from 25 to 50 to 100 units H_2_O_2_, pink). (**D**) Mean expression levels for model (left) and experiments (right) of pulsatile gene *PkatG* for step (orange) or gradual H_2_O_2_ treatments (from 25 µM to 50 µM to 100 µM H_2_O_2_, pink) as depicted in panel C (n≥2 repeats). (**E**) Snapshot of *PgrxA* expression at steady state for step (left and middle) or gradually (right, as shown in panel C-D) treated wild-type (left) cells and Δ*katG* cells (middle and right) after 120 min of treatment with 100 µM H_2_O_2_ (Scale bar = 10 µm). (**F**) Gradual treatment enables adaptation of Δ*katG* cells. Mean elongation rate for Δ*katG* cells (located at a position with 3 barrier cells) treated with 100 µM H_2_O_2_ in a step (orange) or gradual (pink) manner (as depicted in panel C-E, n≥2 repeats). (**G**) Boxplots indicate the median of time taken to reach the peak gene expression for individual mother cells under 100 µM H_2_O_2_ treatment (gradual: blue, pulsatile: pink; box length extends to the lower and upper quartile with error bars representing the range, n≥3 repeats per gene) (**H**) Mean coefficient of variation (C.V.) for peak gene expression values of frontier cells showing cell-to-cell heterogeneity under 100 µM H_2_O_2_ treatment (blue: gradual, pink: pulsatile genes, error bars represent standard deviation, n≥3 repeats per gene).

These analyses suggest that pulsatile genes are important in response to sudden stress, whereas gradually induced genes are more important during prolonged stress. Consistent with this hypothesis, deletion of the pulsatile *katG* gene makes cells extremely sensitive to sudden H_2_O_2_ treatment, but these cells were still able to survive a gradually increasing dose of H_2_O_2_ reaching the same final concentration [Figure 5E, F, Movie S4]. That is, the expression pulse of *katG* appears to be critical for the rapid production of catalase enzymes to counteract the transient spike in intracellular H_2_O_2_ but *katG* expression becomes dispensable after the initial adaptation delay, as noted before^62^. H_2_O_2_ tolerance during prolonged exposure is then provided by the gradually upregulated AhpCF alkyl hydroperoxidase, which scavenges lower concentrations of H_2_O_2_ efficiently^30^.

### Pulsatile genes show higher cell-to-cell expression heterogeneity

We next asked to what extent the expression of pulsatile and gradually induced genes is coordinated within the same cell. Firstly, we found that the time of induction after the onset of stress was shorter and less variable across cells for the pulsatile genes [Figure 5G].

However, the amplitude of the expression peak reached by individual cells was more variable for pulsatile genes compared to gradually induced genes [Figure 5H, Figure S1, S3]. Regulation of pulsatile genes, therefore, appears to prioritise precise control of the timing over the magnitude of the response [Figure 5G-H, S3]. That is, this form of regulation function to activate a response very quickly, if not accurately, in the face of rapid onset stress.

Next, we constructed a dual-reporter strain with two fluorescently labelled transcriptional reporters: the pulsatile P*katG*-YFP reporter expressed from a plasmid and the gradually induced P*grxA*-CFP reporter inserted on the chromosome [Figure 6A, Movie S5]. We also constructed a control strain with two P*grxA* reporters, one marked with CFP on the chromosome and one marked with YFP on the plasmid, to account for variability due to differences in fluorescent proteins (YFP versus CFP) and the difference in gene copy numbers of the reporters (chromosomal versus multi-copy plasmid) [Figure 6A, Movie S6]. The mean expression dynamics of the dual reporter strains matched our previous analysis for P*katG* and P*grxA* in single reporter strains [Figure 6B, Movie S5]. Interestingly, for the control strain, the plasmid-expressed P*grxA* displayed a slightly higher expression pulse compared to the P*grxA* reporter on the chromosome, even after normalising for differences in the steady-state level [Figure 6C]. This is a subtle but significant effect and explained by our model, which showed that the plasmid copy number increases transiently when the cell growth rate slows during the onset of H_2_O_2_ treatment [Figure 6C]. This effective increase in gene dosage leads to an additional expression boost for the plasmid-based promoter.

**Figure 6:**
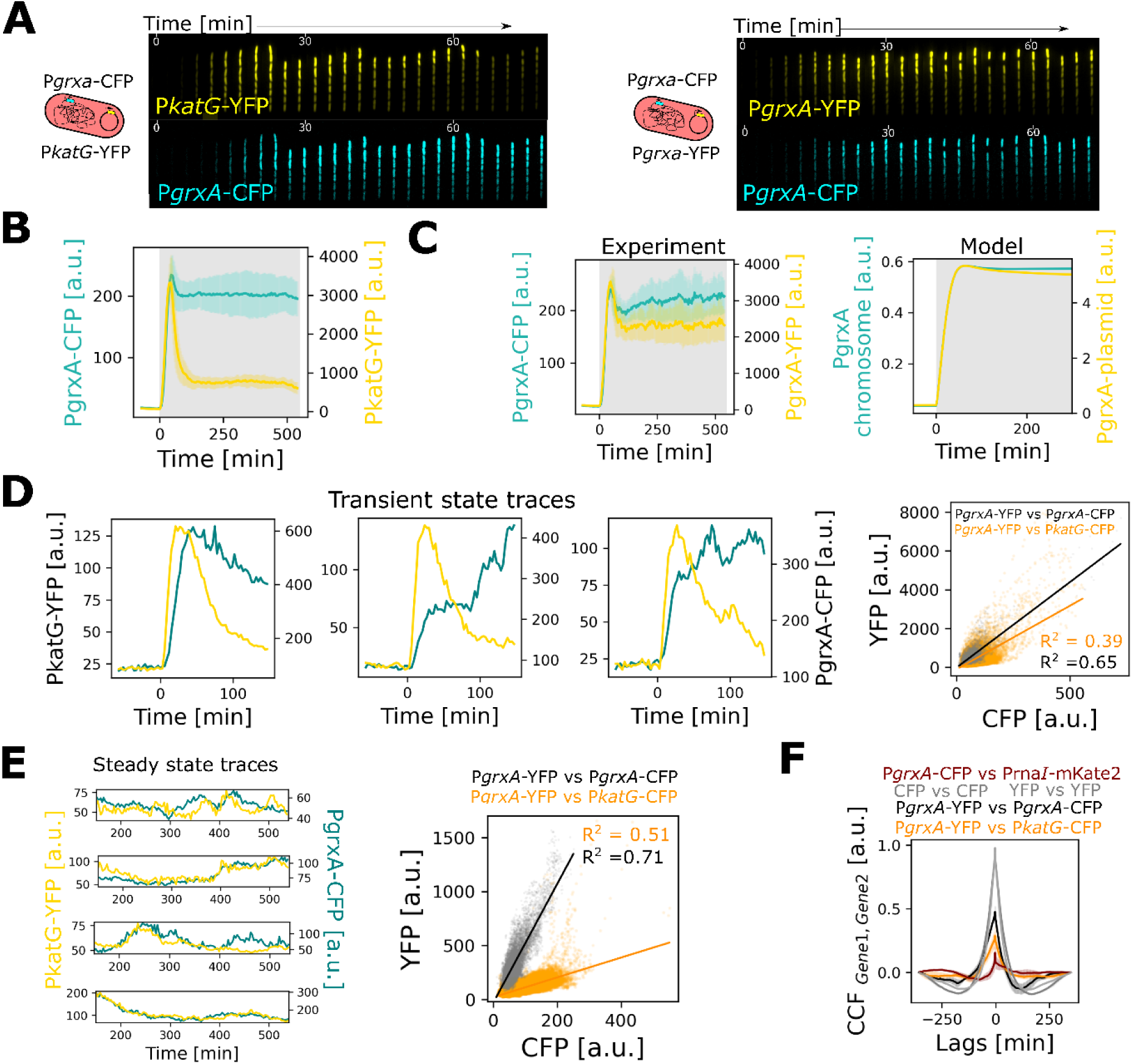
Coordination of pulsatile and gradual gene regulation in single cells. (**A**) *E. coli* dual reporter strains with P*katG*-YFP on plasmid + P*grxA*-CFP on chromosome (left) and P*grxA*-YFP on plasmid + P*grxA*-CFP on chromosome (right, control strain). Kymographs represent gene expression levels for both reporters in cells treated with 100 µM H_2_O_2_ at t=0 minutes. (**B**) Mean expression of P*grxA*-CFP (cyan) and P*katG*-YFP (yellow) for frontier cells under 100 µM H_2_O_2_ provided at t = 0 min (n= 3 repeats, error bars: standard deviation). (**C**) Mean expression of P*grxA*-CFP (cyan) and P*grxA*-YFP (yellow) for frontier cells under 100 µM H_2_O_2_ provided at t = 0 min from experiments (left) and model output (right) (n= 3 repeat, error bars: standard deviation). (**D**) (left) Representative expression of P*katG*-YFP (yellow) and P*grxA*-CFP (cyan) for individual mother cells after treatment with 100 μM H_2_O_2_ from t = 0 min. (right) Peak expression of P*grxA*-YFP vs. P*grxA*-CFP (black) and P*grxA*-YFP vs. P*katG*-CFP (orange) for mother cells after treatment with 100 μM H_2_O_2_. The lines represent linear fits. (**E**) (left) Representative expression traces of P*katG*-YFP (yellow) and P*grxA*-CFP (teal) for individual mother cells during steady state with 100 μM H_2_O_2_ treatment from t = 0 min. (right) P*grxA*-YFP vs. P*grxA*-CFP (black) and P*grxA*-YFP vs. P*katG*-CFP (orange) expression for mother cells during steady-state with 100 μM H_2_O_2_ treatment. The lines represent the linear regression fit to the data sets. (**F**) Mean temporal cross-correlation of steady-state expression dynamics between P*grxA*-YFP and P*rna1*-mKate2 (red), P*grxA*-YFP and P*grxA*-YFP or P*grxA*-CFP and P*grxA*-CFP (dashed grey), P*grxA*-YFP and P*grxA*-CFP (black), P*grxA*-YFP and P*katG*-CFP (orange) for mother cells under 100 μM H_2_O_2_ treatment.

Focusing on the dual reporter strain, we found that P*katG* and P*grxA* expression in the same cell showed relatively little correlation during the transient expression pulse [Figure 6D, Movie S5]. However, both promoters exhibited substantial fluctuations in gene expression during prolonged treatment, and we found that these fluctuations were closely correlated [Figure 6E, Movie S5]. This was evident from gene expression levels at discrete time points and from temporal cross-correlation analysis [Figure 6F]. Together, our single-cell analysis revealed that pulsatile and gradually induced genes are tightly coordinated during prolonged stress, but appear to be uncoupled in regulation during the transient response to sudden stress.

### Differentially-regulated genes show distinct spatial patterns in cell populations

Spatial patterns of gene expression are important in the oxidative stress response because of the ability of cells nearest the stress to protect cells that are further away. This phenomenon is remarkably effective, with each individual cell reducing the local H_2_O_2_ concentration by around 30% in its vicinity inside the one-dimensional microfluidic channel^26^. Within colonies, cells are protected in all three dimensions by surrounding cells leading to even steeper spatial H_2_O_2_ gradients from the edge to the interior of the population^28^. However, how this spatial structuring affects the whole oxidative stress regulon is unknown. We, therefore, asked whether the different temporal expression patterns we observe across the regulon were associated with different spatial expression patterns.

Quantifying expression levels in the channels of the microfluidic device revealed clear differences in the spatial patterning of regulation across genes. Downregulated genes exhibited an inverse gradient to that of the stress, with the lowest expression seen in the frontier cells that are closest to the H_2_O_2_ source [Figure 7A, C, Movie S2]. Even passively induced genes that are not part of the OxyR regulon showed a spatial gradient in expression level that was driven by a more pronounced inhibition of growth for cells that are located closer to the H_2_O_2_ source [Figure 7C]. Turning to upregulated genes, we see a steeper decay in expression in space as one moves away from the source of stress for pulsatile genes as compared to the gradually induced genes, in both our model and experiments [Figure 7, Movie S1]. That is, pulsatile genes are strongly upregulated in relatively few cells that are close to the H_2_O_2_ source, which generates significant cross protection, such that only the most stressed cells show expression. The gradually upregulated genes then activate more evenly in larger numbers of cells to provide lasting protection.

**Figure 7:**
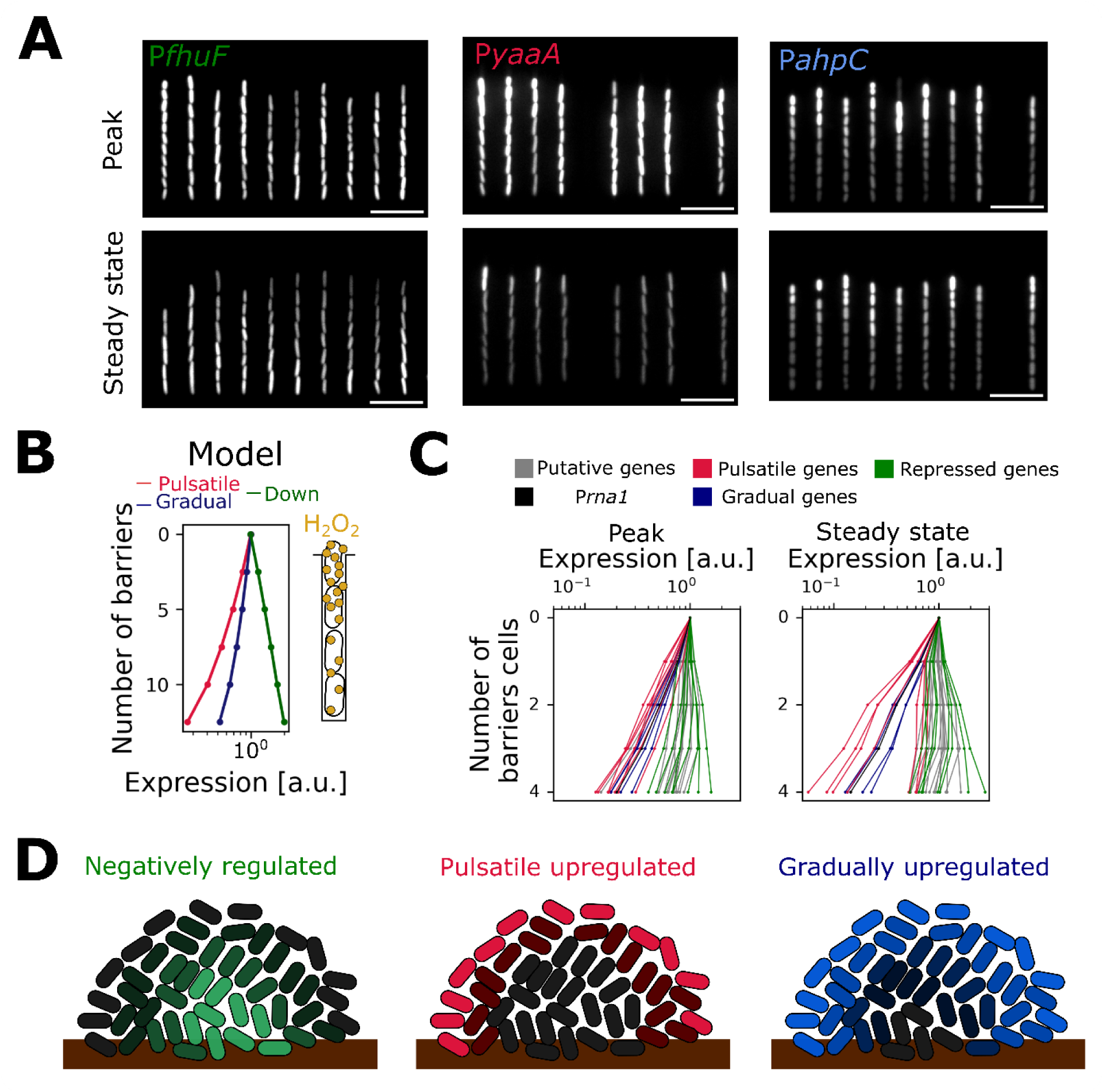
Spatio-temporal expression patterns across the OxyR regulon. (**A**) Snapshots of cells with representative reporters for the three categories of gene regulation: P*fhuF* (downregulated), P*yaaA* (pulsatile) and P*ahpC* (gradually induced) during peak expression (top) and at steady-state (bottom) with 100 µM H_2_O_2_ treatment. (**B**) Scavenging of H_2_O_2_ by bacteria creates H_2_O_2external_ gradients in the growth trench from the source of treatment at the open end to the mother cell at the closed end. Model output for the expression of pulsatile (pink), gradually induced (blue) and downregulated (green) genes for cells across the growth trench (increasing number of barrier cells) relative to the expression of frontier cells. (**C**) Mean expression of 31 transcriptional reporters for cells with different number of barrier cells with 100 µM H_2_O_2_ treatment relative to the expression of frontier cells during peak expression (left) and steady-state (right) (pulsatile: pink, gradual: blue, down: green, constitutive P*rna1*: black, putative: grey; n≥3 repeats per gene). (**D**) Schematic depicting the spatial variation across a bacterial population in the expression of OxyR-controlled genes that are downregulated (left), pulsatile induced (center) and gradually induced (right) under H_2_O_2_ treatment.

## Discussion

The rapid and coordinated regulation of stress response genes is critical for survival in a changing environment^74^. However, bulk measurements provide limited insights into how gene expression dynamics are coordinated in space and time within cell populations. To address this, we leveraged the power of single-cell transcriptional reporters to follow the bacterial oxidative stress response under continuous H_2_O_2_ treatment. Our work revealed a diverse set of dynamics across 31 oxidative stress response genes. Despite this diversity, our modelling and experiments show that OxyR binding dynamics together with the effects of stress on the cell growth rate can explain the different gene regulation patterns. Promoters with a negative *K_ind_* exhibit reduced expression rate, despite the fact that protein levels transiently increase due to a slow-down of cell growth. For upregulated genes, promoters with a high *K_D_* drive a strong transient expression pulse after H_2_O_2_ treatment leading to pulsatile expression, whilst promoters with a lower *K_D_* activate genes more slowly and sustain elevated expression rates throughout prolonged treatment. Moreover, by using a dual reporter, we show that the expression of different genes in the stress response diverges during the initial rapid response to treatment but then becomes closely correlated at steady-state.

The dynamics we observe are broadly consistent with previous bulk measurements^4,36,49^ for time points shortly after the start of treatment [Table 1]. We observe differences at later time points [Figure S4]. However, these differences are to be expected as bulk methods typically study gene expression in dense cultures after a single high dose of H_2_O_2_. H_2_O_2_ levels decay quickly in bulk cultures and the stress response abates, which is not the case with continuous supply of H_2_O_2_ in microfluidic chips^4,37,50^.

What is the evolutionary function of the different patterns in gene regulation that we observe? Our model and experiments suggest that the higher H_2_O_2_ sensitivity of pulsatile genes provides a rapid defence against sudden bursts of intracellular H_2_O_2_. Interestingly, several of the pulsatile genes including *yaaA*^38^, *clpS*^37^ and *hemH*^41^ are involved in iron regulation. Their pulsatile induction is likely beneficial to counter the rapid lethality of H_2_O_2_ experienced due to the Fenton reaction^62,75–78^. Being more sensitive to local H_2_O_2_ fluctuations, pulsatile genes therefore display high cell-cell variability in expression magnitude and steeper spatial gradients compared to gradually induced genes. Of the two H_2_O_2_ scavenging enzymes, pulsatile catalase (*katG*) is important to protect against immediate stress experienced by cells on the periphery of a population while gradually induced alkyl hydroperoxidase (*ahpCF*) efficiently scavenges low H_2_O_2_ levels that reach the rest of the population. Gradually induced thioredoxins (grxA^78^, trxC^46^) regulate the redox status of proteins, including OxyR. These genes, therefore, show a more consistent response both in space and time, with more cells contributing to protection at the steady-state. In this way, the action of a single transcription factor is able to orchestrate the stress response to generate both individual and collective protection.

## Acknowledgments

We thank Somenath Bakshi, Georgia Isom and members of the Uphoff and Foster labs for their discussions and comments on the manuscript.

## Author contributions

Conceptualization and design of study: DC, KRF, SU; Data collection: DC; Development of the computational model: DC, SU; Data analysis and interpretation: DC, KRF and SU; Writing of the article: DC, KRF and SU; Supervision: KRF, SU

## Materials availability

Further information and requests for resources and reagents should be directed to and will be fulfilled by Stephan Uphoff and Kevin Foster.

## Data and code availability

All the raw data collected for analysis in this study along with the custom-built python codes for analysis and simulations will be freely and openly available on the Oxford Research Archive. Any further information about data and code is available upon request by the corresponding authors.

## Supplementary movies

**Movie S1:** Movie of P*trxC*-CFP (gradually induced, top) and P*hemH*-CFP (pulsatile gene, bottom) fluorescence for cells growing in microfluidic trenches under 100 µM H_2_O_2_ treatment starting at time = 0 min. (Scale bar: 10µm)

**Movie S2:** Movie of P*fhuF*-CFP fluorescence for cells growing in microfluidic trenches under 100 µM H_2_O_2_ treatment starting at time = 0 min. (Scale bar: 10µm)

**Movie S3:** Movie of P*rna1*-CFP fluorescence for cells growing in microfluidic trenches under 100 µM H_2_O_2_ treatment starting at time = 0 min. (Scale bar: 10µm)

**Movie S4:** Movie of P*grxA*-CFP fluorescence for Δ*katG* cells growing in microfluidic trenches under 100 µM H_2_O_2_ treatment applied gradually (top) or suddenly (bottom) starting at time = 0 min. (Scale bar: 10µm)

**Movie S5:** Movie of P*katG*-mYPet (top) and P*grxA*-CFP (bottom) fluorescence for dual reporter strain growing in microfluidic trenches under 100 µM H_2_O_2_ treatment starting at time = 0 min. (Scale bar: 10µm)

**Movie S6:** Movie of P*grxA*-mYPet (top) and P*grxA*-CFP (bottom) fluorescence for dual reporter strain growing in microfluidic trenches under 100 µM H_2_O_2_ treatment starting at time = 0 min. (Scale bar: 10µm)

## Materials and Methods

### Strains and plasmids

We used strains derived from *E. coli* K12 AB1157 for our experiments. All the strains had a constitutively expressed P*rna1*-mKate2 marker aiding the segmentation of cells during microscopy and *flhD* deletion to stop the cells from escaping out of the microfluidic growth channels during time-lapse microscopy. The reporter plasmids for genes used in this study were obtained from a transcriptional reporter library of PSC101 plasmids ^83^. The reporter plasmids comprised of GFPmut2 fluorescence protein after the promoter region of a given gene or operon and had a kanamycin resistance marker. GFPmut2 was replaced with SCFP3A fluorescent protein for P*katG*, P*ahpC*, and P*grxA* using Gibson Assembly (NEB)^28^. The reporter plasmids were mini-prepped and transformed in our AB1157 background strain. Strains were selected on 25 μg/mL kanamycin resistance LB agarose plates and checked for fluorescence signal by microscopy snapshots.

The chromosomal reporter for P*grxA*-SCFP3A was constructed by inserting P*grxA*-SCFP3A-kanamycin using λ-red recombination in endogenous loci (*aslA*) on the chromosome of DH5α strain expressing pKD46-ampicillin. The successful colonies were selected by growing them overnight at 37°C on kanamycin plates and subsequently re-streaking again on kanamycin selective plates to remove the temperature sensitive pKD46 plasmid. Then, the insert was moved into our strain of interest that had P*rna1*-mKate2 marker and *flhD* deletion, using P1-phage transduction. The successful colonies were selected on kanamycin plates and tested for the insert using colony PCR, after which, the colonies were restreaked and grown overnight thrice to remove the phage. Colony PCR was again performed before storing the strain at -80°C. P*grxA*-mYPet and P*katG*-mYPet plasmids with ampicillin resistance marker were obtained by genetically editing P*grxA*-SCFP3A-kanamycin and P*katG*-SCFP3A-kanamycin plasmids using Gibson assembly. We first replaced CFP with mYPet fluorescent protein and the resultant plasmid was edited again using Gibson assembly to replace kanamycin with an ampicillin resistance marker. The final strain was checked by colony PCR, microscopy snapshot, and re-streaking on 100 μg/mL ampicillin plates. Finally, the P*grxA*-mYPet-ampicillin and P*katG*-mYPet-ampicillin plasmids were transformed in the P*grxA*-SCFP3A-kanamycin chromosomal fluorescent reporter strain to construct the dual reporter strain. The strain was imaged to test for dual fluorescence using microscopy snapshots and selected on plates supplemented with 25 μg/mL kanamycin and 100 μg/mL ampicillin.

### Media and growth conditions

Strains were stored in 20% glycerol stocks at -80°C. Strains were streaked on freshly prepared LB agar plates supplemented with appropriate antibiotics for selection (25 μg/mL kanamycin and/or 100 μg/mL ampicillin) and incubated at 37°C. One colony was picked from the overnight plates and added in minimal M9 media with appropriate antibiotics for overnight growth at 37°C in a shaking incubator at 200rpm. The minimal media was prepared by mixing M9 salts (15 g/L KH_2_PO_4_, 64 g/L Na_2_HPO_4_, 2.5 g/L NaCl, and 5.0 g/L NH_4_Cl), 2 mM MgSO_4_, 0.1 mM CaCl_2_, 0.5 mg/mL thiamine, MEM amino acids, 0.1 mg/mL L-proline, and 0.2% glucose. The next day, cells were diluted 1:50 in minimal M9 media and grown for ∼3 hours for the microfluidic experiments. 0.85 mg/mL Pluronic F127 was also added to this culture and minimal media to avoid cell aggregation when loading the cells on a microfluidic chip. Minimal media with or without H_2_O_2_ was flown continuously through microfluidic chips using syringe pumps. The concentration of H_2_O_2_ used was as specified in the figure legends or text, and was added to the minimal M9 media just before setting up the experiments.

### Microfluidic chip preparation and setup

Microfluidic setup and preparation of ‘mother machine’ devices were performed as described in Choudhary *et al*^28^.

### Time-lapse microscopy

Time-lapse imaging was performed using a Nikon Ti-E inverted fluorescence microscope equipped with 100x NA 1.40 immersion oil objective, motorized stage, sCMOS camera (Hamamatsu Flash 4), LED excitation source (Lumencor SpectraX), and operated with a perfect focus system. Exposure times were 100 ms for P*rna1*-mKate2 (λ = 555 nm), 300 ms for MutL-mYPet (λ = 508 nm), 75ms for GFPmut2 reporter (λ =470 nm) and 75 ms for sCFP3 reporter (λ = 440 nm) using 50% of maximal LED excitation intensities. The excitation and emission lights were separated using a triband dichroic and individual emission filters. The microscope chamber (Okolabs) was maintained at 37°C throughout the experiments and images were captured every 3 min.

### Mother machine data processing and analysis

The microscopy time-lapse .nd2 files for the experiments were processed using BACMMAN plugin^84^ in Fiji^85^, which were subsequently analysed using custom made Python scripts. BACMMAN first performs pre-processing which corrects for drifts during imaging and aligns growth channels spatially over time. To obtain cellular parameters, we used the P*rna1*-mKate2 marker to segment and mark the cell outlines against the background and then used this as a mask for other fluorescence channels. In all the fluorescence channels (CFP, GFP and mYPet), the mean fluorescence intensity inside the mask of each cell mask were computed. The cells were tracked over time using the segmentation masks of mKate2 channel to provide cell lineage information. BACMMAN generates output in separate excel files containing cell growth characteristic, P*rna1*-mKate2 intensity data and one for each fluorescence data. These files were then further analyzed using a custom python pipeline to compute cell parameters as described in the following section.

### Cell parameter calculations

#### Elongation Rate

The instantaneous elongation rate at time *t* was calculated based on the log-difference in cell length *L*_*t*_ at time between consecutive frames as 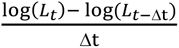. For calculating the elongation rates of cells at different positions in the growth trench, cells were tracked according to their initial position until the number of barrier cells decreased by 2.

#### Gene expression

Reporter fluorescence intensity values were averaged over the area of each cell.

#### Number of barrier cells

The number of cells that are located between the open end of a trench and the cell being analysed.

#### Promoter activity

The instantaneous promoter activity at time *t* was calculated as the cell averaged rate of change of total fluorescence intensity of a cell. It was computed as^16^: 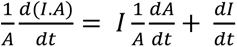 where *I* is the mean fluorescent intensity of cell and A is the cell area.

Here, 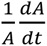 was given by the instantaneous elongation rate of the cell. For calculating the promoter activity values of cells at different positions in the growth trench, cells were tracked according to their initial position until the number of barrier cells decreased by 2.

#### Expression Rate

The instantaneous expression rate at time *t* was calculated based on the difference in cell intensity *I*_*t*_ at time between consecutive frames as 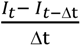. For calculating the expression rates of cells at different positions in the growth trench, cells were tracked according to their initial position until the number of barrier cells decreased by 2.

#### Peaky expression

90^th^ percentile of the mean fluorescence intensities from 12 to 90 minutes post H_2_O_2_ treatment.

#### Steady expression

Average of the mean fluorescence intensities from 150 minutes post H_2_O_2_ treatment.

#### Time to peak expression for single cells

Time in minutes post treatment until 100 minutes when individual cells show maximum fluorescence intensity.

#### Peak expression for single cells

Mean fluorescence intensity for individual cells from 18 minutes until 90 minutes post treatment. This value was subtracted by the mean fluorescence intensity without H_2_O_2_ treatment i.e. basal fluorescence intensity.

#### Coefficient of variation

The CV values were calculated as the standard deviation divided by the mean.

#### Relative expression in spatial dimension

Expression intensity of cells with atleast 1 barrier cells were divided by the intensity of outermost cell (i.e. cell with no barrier cells) for each growth channel at a given time point.

#### Cross-correlation analysis

The temporal cross-correlation between the different reporter intensity traces of mother cells was computed using the statsmodel library in Python. Correlation values from individual growth trenches were then averaged over all observed growth trenches.

### Linear regression analysis

The linear regression analysis was performed in Python using the stats.linregress function in scipy library which computes the least square regression for a linear fit between two sets of data points.

### Oxidative stress response model

We modelled the oxidative stress response model as shown in Figure 4A and previously described in Choudhary *et al*^71^. Here, external H_2_O_2_ is provided at the rate *R*_*influx*_. Intracellular concentration of H_2_O_2_ ([*H*_2_*O*_2_]_*cell*_) oxidises OxyR with rate *K*_*ox*_and converts it into oxidised form that induces multiple genes in its regulon. For simplicity, we modelled the induction of important genes of stress response regulon by OxyR: glutaredoxin-1 (grxA) that converts OxyR oxidised back to its reduced form with rate *K*_*red*_, scavengers: alkyl hydroperoxidase (ahpC) and catalase (katG) that reduce [*H*_2_*O*_2_]_*cell*_and a reporter gene that does not directly or indirectly affect the OxyR regulation. All the genes are produced at a basal expression rate (*R_grxA,basal_*, *R_katG,basal_*, *R_ahpC,basal_*, *R_reporter,basal_*) and their expression is modulated upon induction by OxyR. The induction by OxyR is modelled as Michaelis Menten kinetics with maximal induction rate given by *K_ind_* (*K*_*grxA,ind*_, *K*_*ahpC,ind*_, *K*_*katG,ind*_, *K*_*reporter,ind*_) and half-maximal induction *K_m_* is given by dissociation constant of OxyR from the promoter regions of the genes (*K_D_*) (*K*_*D,grxA*_, *K*_*D,katG*_, *K*_*D,ahpC*_, *K*_*D,reporter*_). Finally, the gene expression reduces due to growth dependent dilution effect where growth rate (*g*) is a function of [*H*_2_*O*_2_]_*cell*_.

This leads to the following equations:

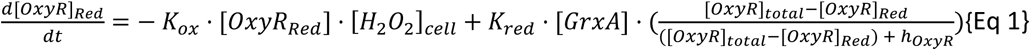

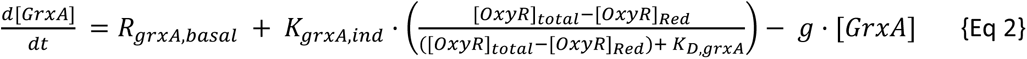

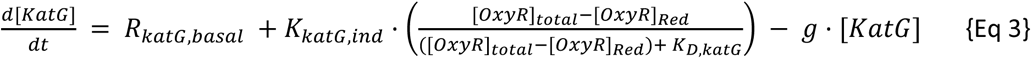

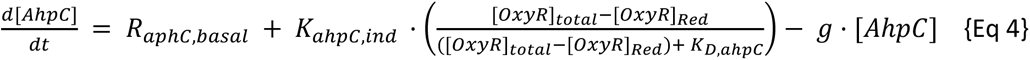

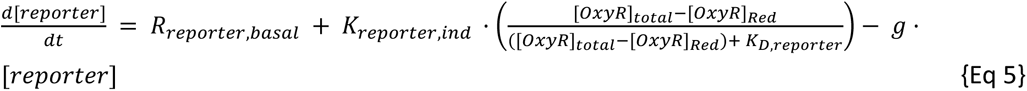

Here *K*_*ind*_ > 0 for positive regulation and *K*_*ind*_ < 0 for negative regulation.

A constitutive gene, that is not part of the OxyR regulon, was modelled such that it was expressed at rate *R*_*constitutive*_ and diluted with growth:

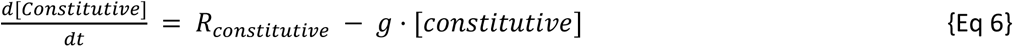

The intracellular [*H*_2_*O*_2_]_*cell*_concentration is determined by the influx of external H_2_O_2_ with rate *R*_*influx*_ · [*H*_2_*O*_2_]_*external*_, a basal endogenous production rate *R*_*H*2*O*2,*basal*_, and scavenging by catalase and peroxidase enzymes with Michaelis-Menten kinetics where *K*_*AhpC*_, *K*_*KatG*_ are the catalytic rate constants and ℎ_*AhpC*_, ℎ_*KatG*_ are the Michaelis constants:

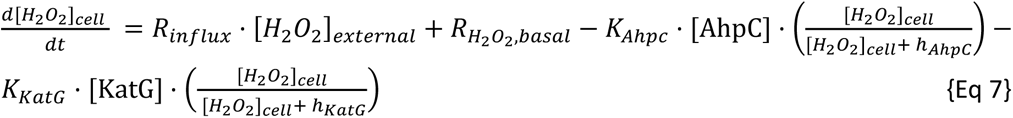

Lastly, the effect of plasmid copy number variation is considered by modifying the equation for reporter gene expression (Eq 5) such that the gene induction is proportional to the plasmid copy number *n*. The change in plasmid copy number is computed as a constitutive expression with rate *R*_*n*_ and diluted by cellular growth as given below.

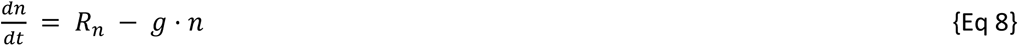

Hence, the variation of reporter gene expression on a plasmid is modelled as follows:

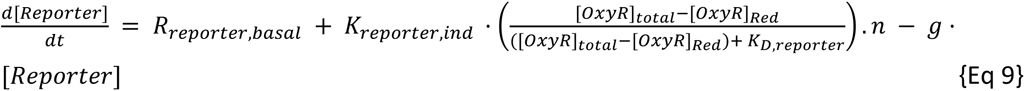

## Key resources table

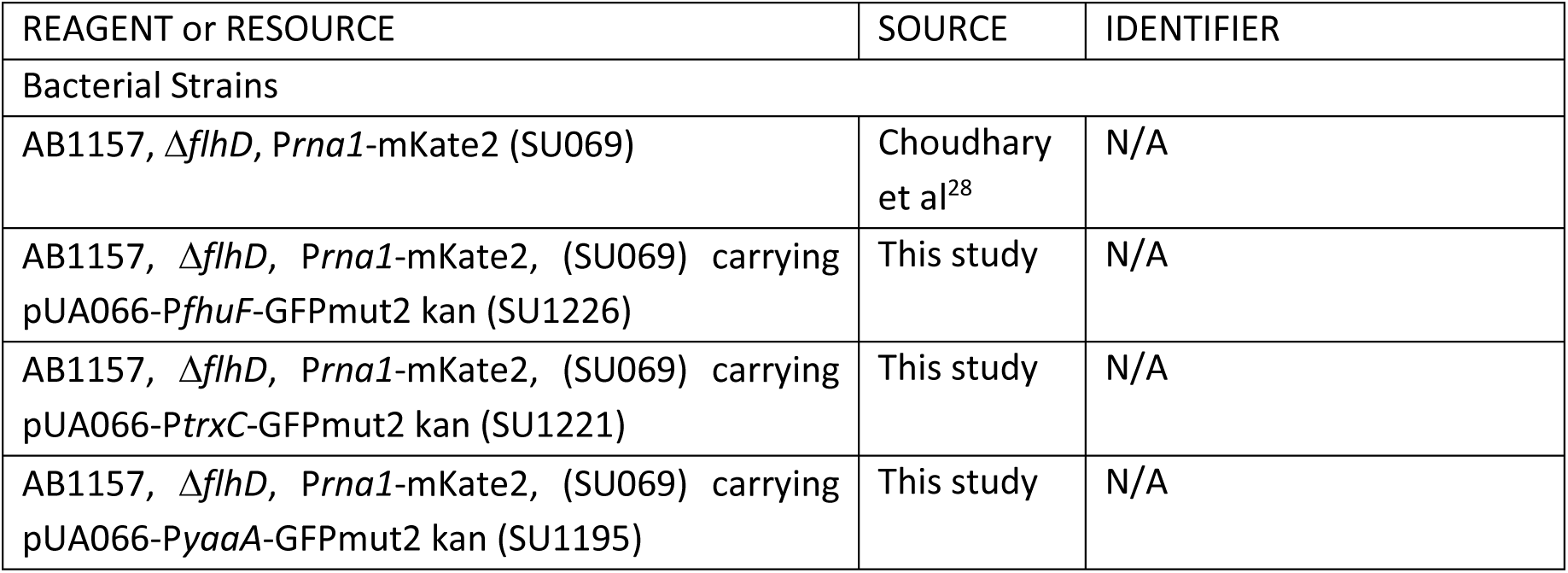
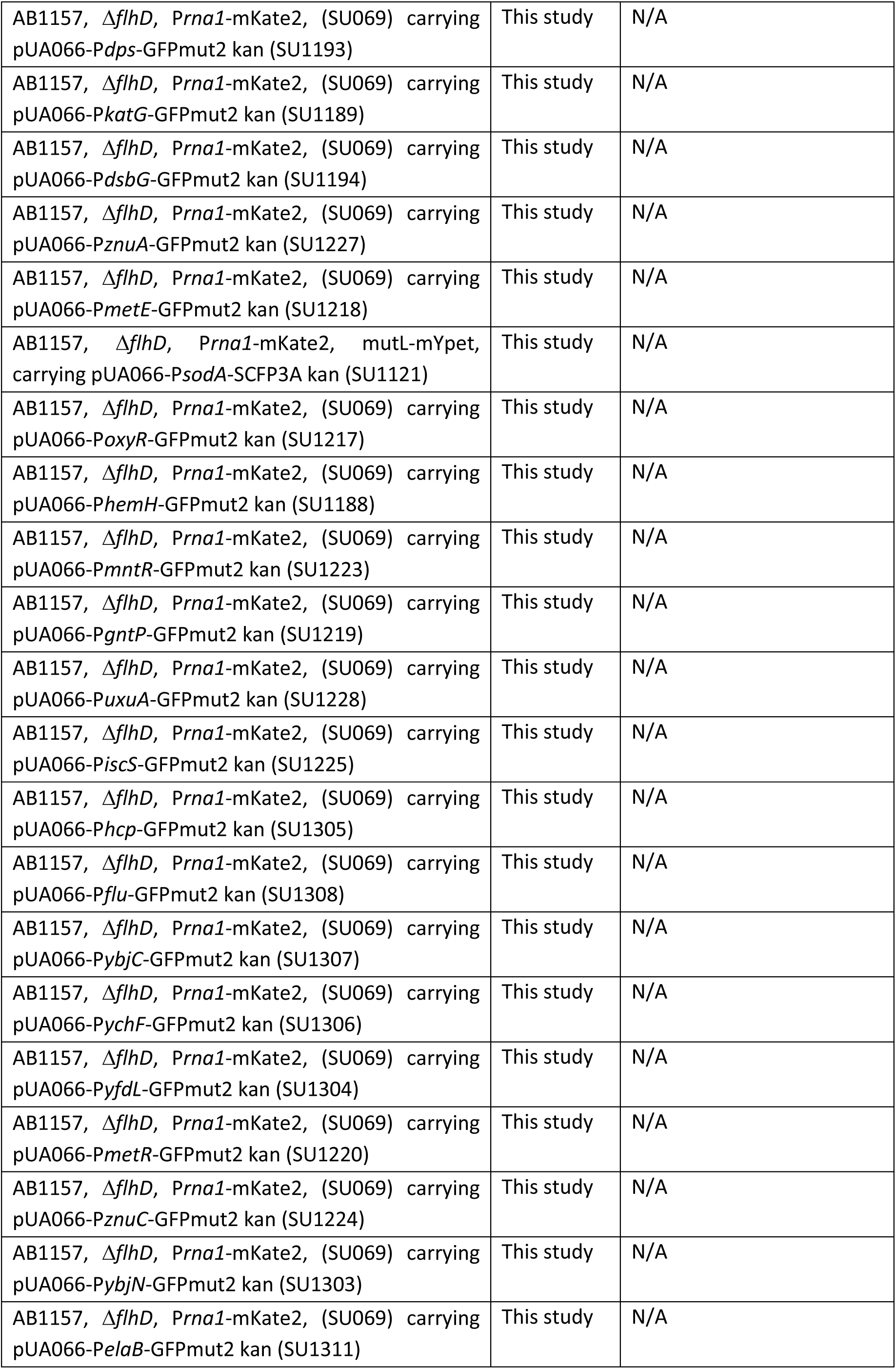
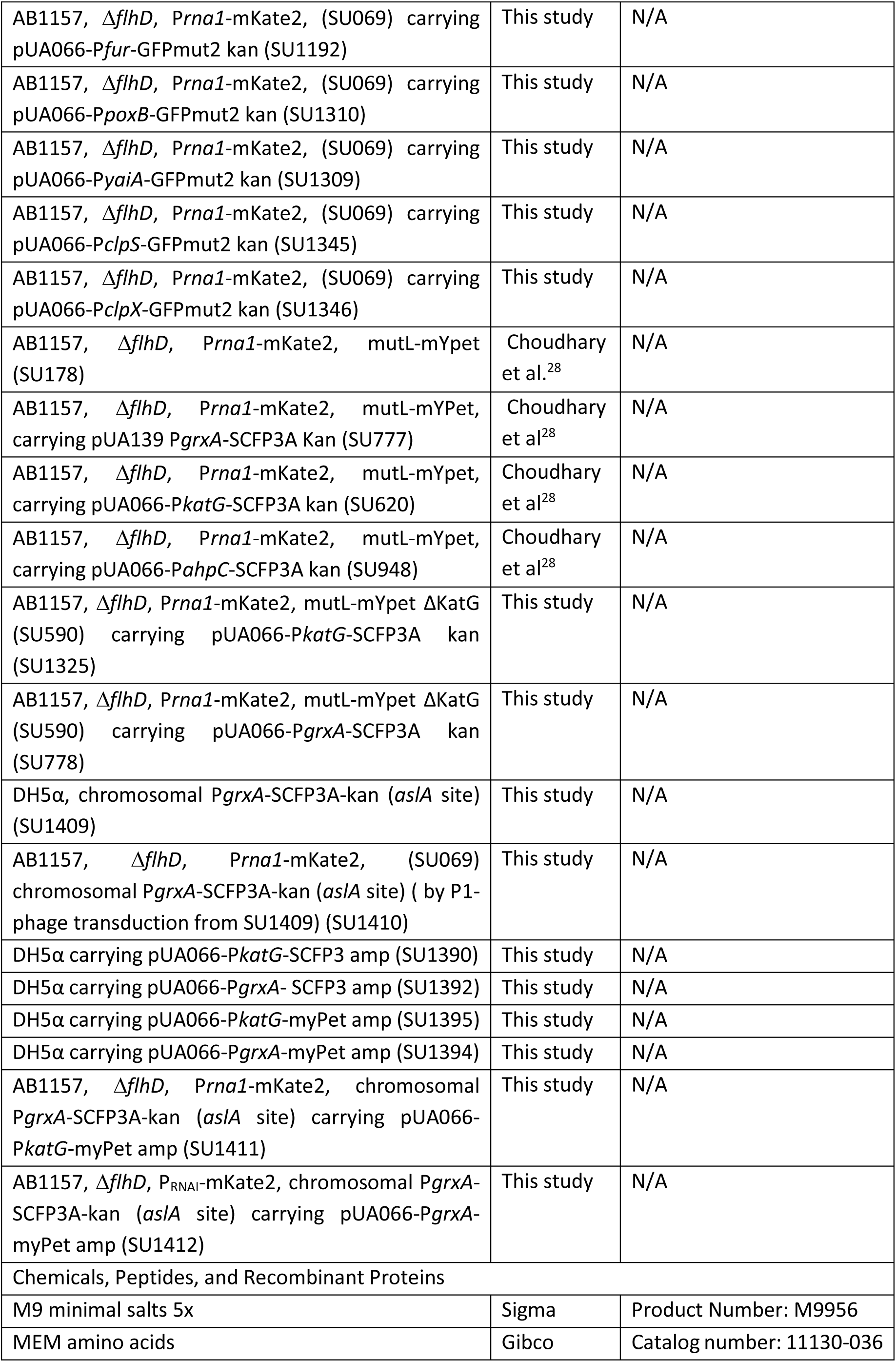
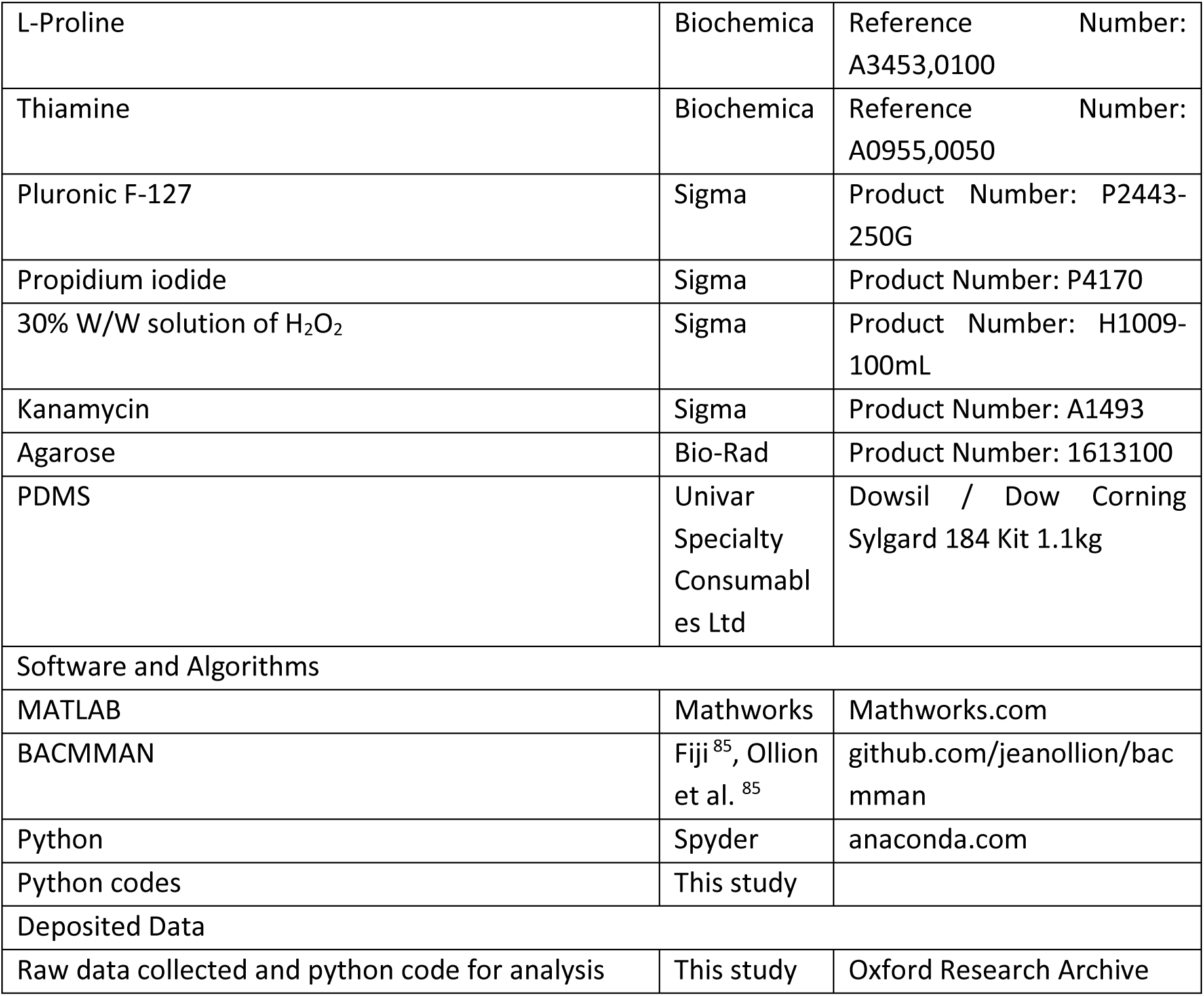

## Supplementary Figures

**Figure S1:**
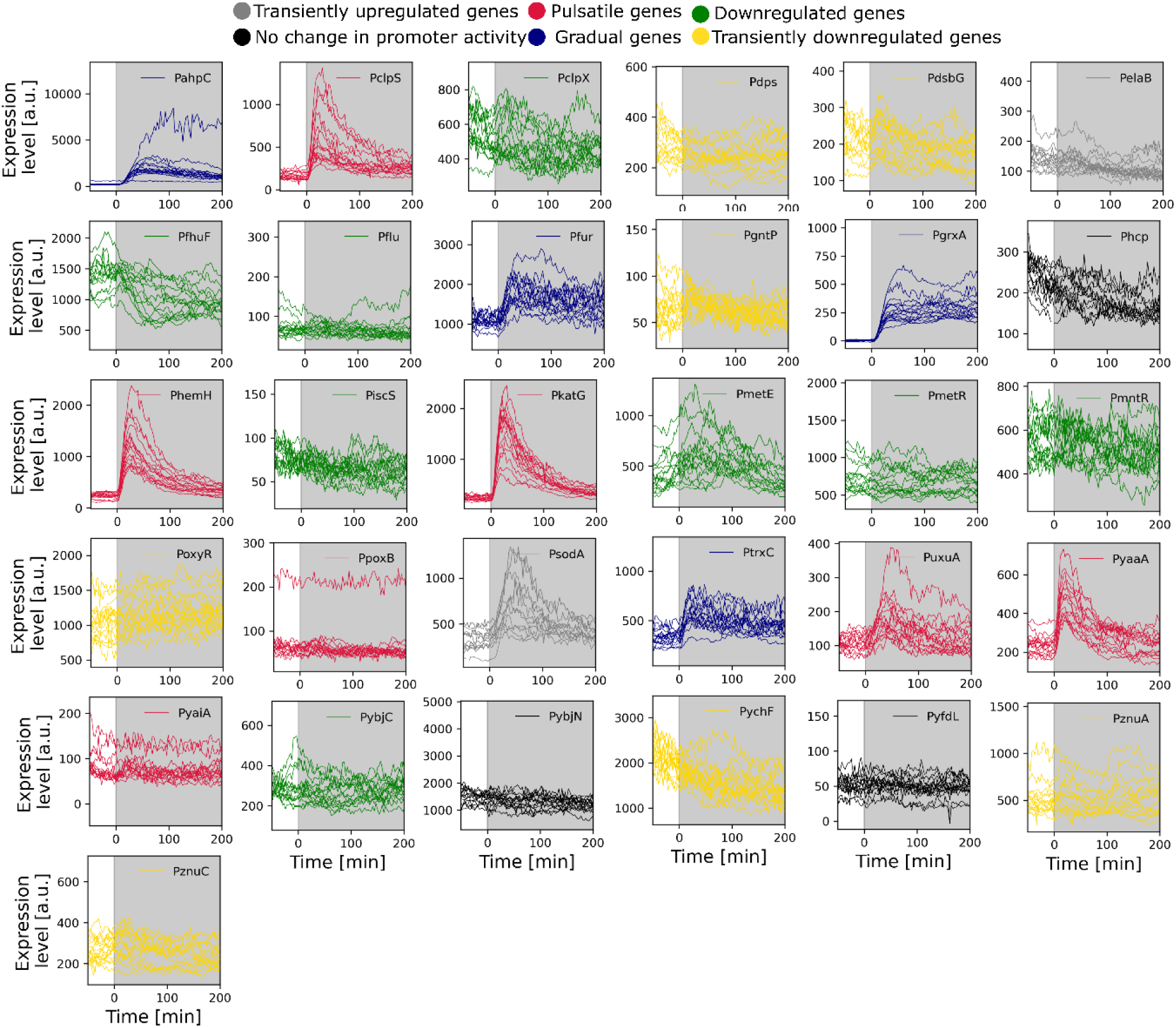
Single cell expression traces of oxidative stress response genes display the cell-cell variability. Single cell expression levels of mother cells for 31 transcriptional reporters with constant 100 µM H_2_O_2_ treatment from t = 0 minutes (shaded region). Line colours correspond to the gene regulation categories (pulsatile: pink, gradual: blue, down-regulated: green, transiently up- (grey) or down- (yellow) regulated, no change in promoter activity: black). n = 15 representative traces per gene.

**Figure S2:**
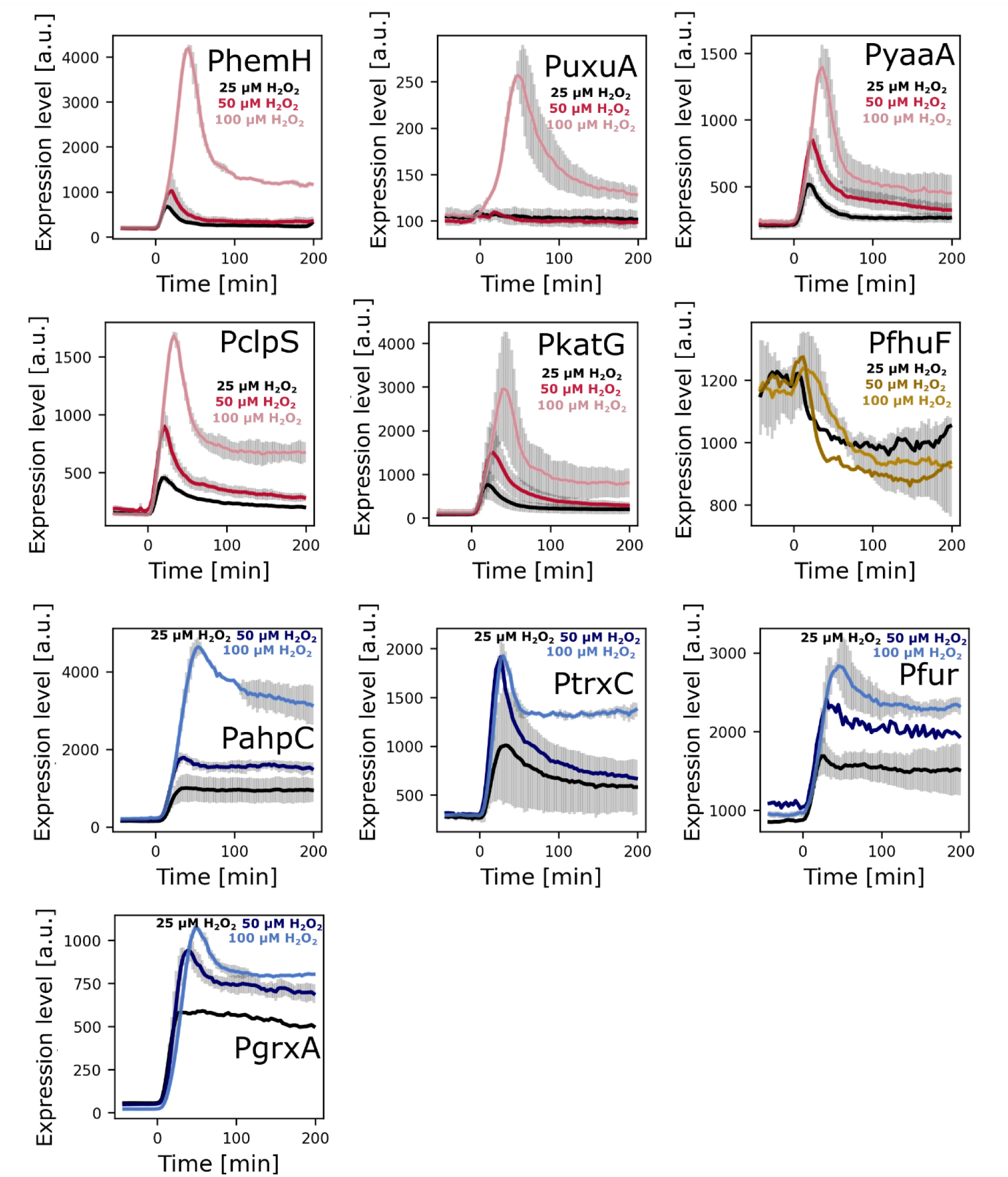
Pulsatile genes exhibit higher expression sensitivity to H_2_O_2_ stress intensity. Mean expression of frontier cells for the indicated transcriptional reporters under 25 µM, 50 µM and 100 µM H_2_O_2_ provided from t = 0 min (red: pulsatile, blue: gradually induced, yellow: downregulated). Error bars: standard deviation (n ≥ 2 repeats per gene).

**Figure S3:**
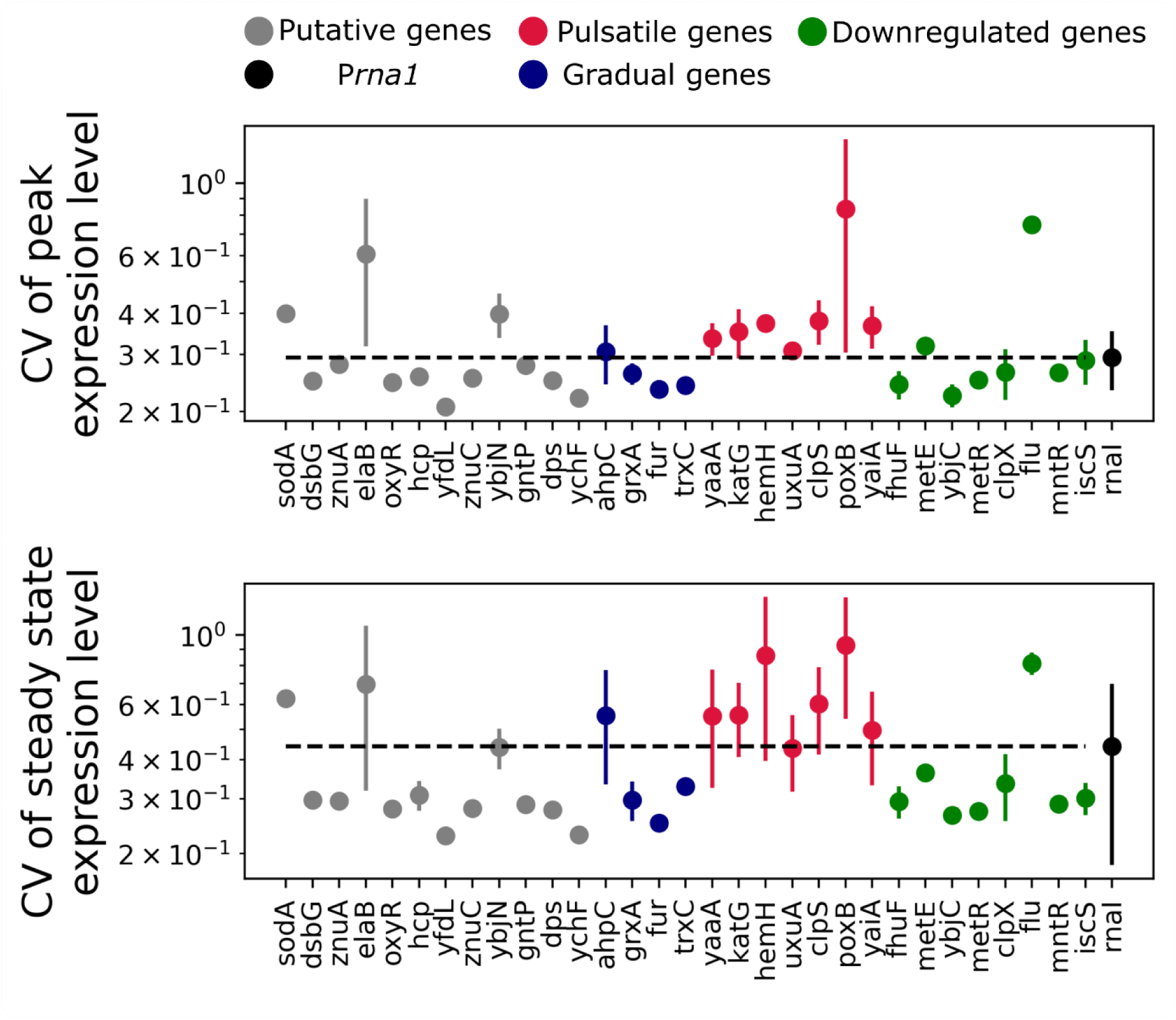
Cell-cell variability in gene expression magnitude for oxidative stress response genes in different regulation categories. Coefficient of variation for peak (top) and steady state (bottom) gene expression values for frontier cells under 100 µM H_2_O_2_ treatment (pulsatile: red, gradual: blue, negative: green, putative: grey, constitutive P*rna1*: black). Error bars represent standard deviation (n ≥ 3 repeats per gene)

**Figure S4:**
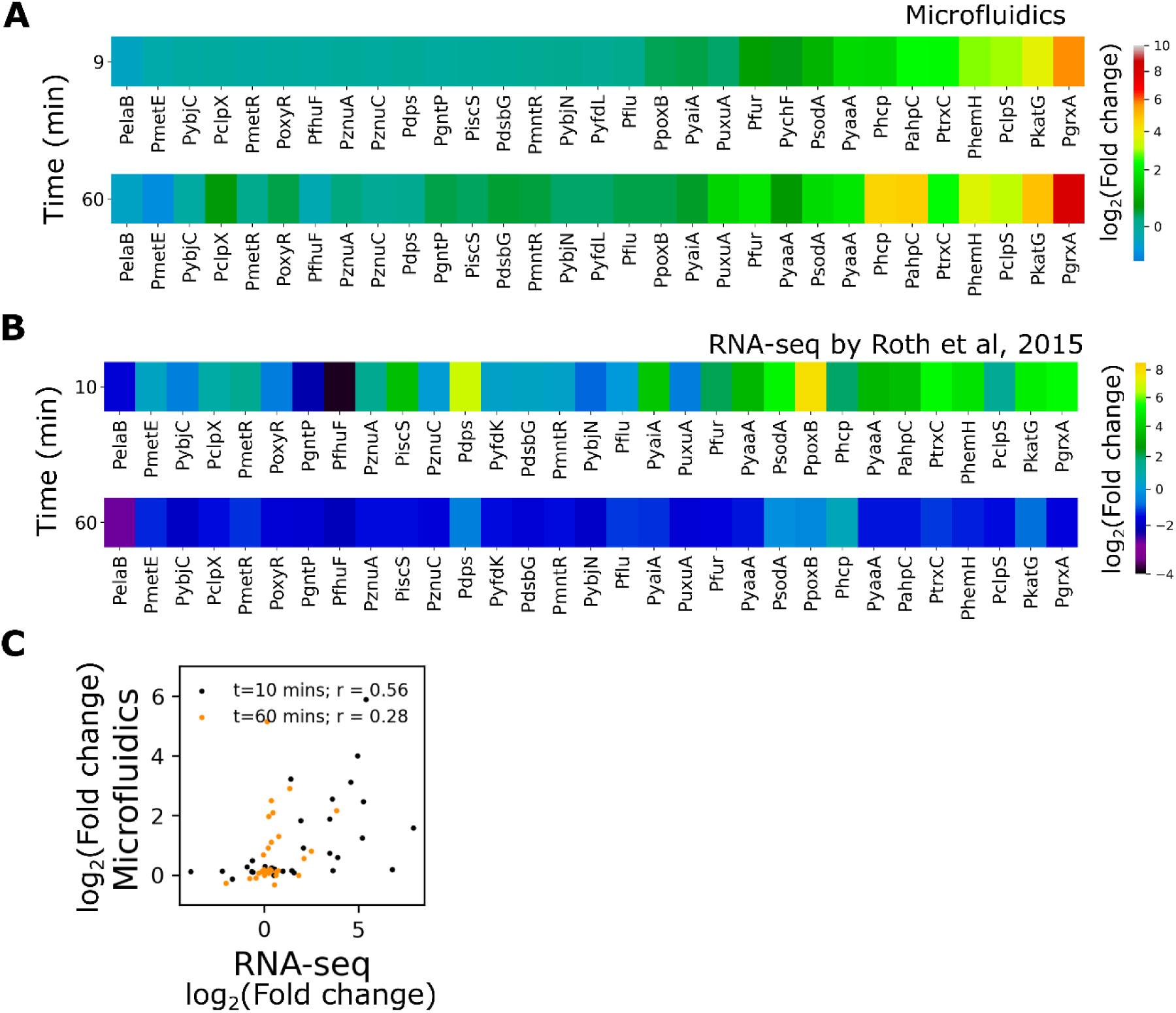
Gene expression level changes in our study compared to the bulk mRNA level changes, reported by Roth *et al*., under H_2_O_2_ stress. (**A**) Heatmap represents mean log_2_-fold change in gene expression relative to basal level of frontier cells for 31 transcriptional reporters at 9 minutes (top) and 60 minutes (bottom) post treatment with 100 µM H_2_O_2_ (n ≥ 1000 cells and ≥2 repeats per gene). (**B**) Heatmap represents data collected by Roth *et al*^4^. showing mean log_2_-fold change in mRNA levels of cells for 31 genes at 10 mins (top) and 60 mins (bottom) after treatment with 2.5 mM H_2_O_2_. (**C**) Comparison of log_2_-fold change in gene expression levels from microfluidic single-cell imaging and mRNA levels in panel A and B at 10 min (black) and 60 minutes (orange) post H_2_O_2_ treatment. Pearson’s coefficient (r) for linear fit between the data sets are indicated in the plot.

